# Embryonically Active Piriform Cortex Neurons Promote Intracortical Recurrent Connectivity during Development

**DOI:** 10.1101/2024.05.08.593265

**Authors:** David C. Wang, Fernando Santos-Valencia, Jun H. Song, Kevin M. Franks, Liqun Luo

**Author notes:** Correspondence (K.M.F.), (L.L.).

## Abstract

Neuronal activity plays a critical role in the maturation of circuits that propagate sensory information into the brain. How widely does early activity regulate circuit maturation across the developing brain? Here, we used Targeted Recombination in Active Populations (TRAP) to perform a brain-wide survey for prenatally active neurons in mice and identified the piriform cortex as an abundantly TRAPed region. Whole-cell recordings in neonatal slices revealed preferential interconnectivity within embryonically TRAPed piriform neurons and their enhanced synaptic connectivity with other piriform neurons. *In vivo* Neuropixels recordings in neonates demonstrated that embryonically TRAPed piriform neurons exhibit broad functional connectivity within piriform and lead spontaneous synchronized population activity during a transient neonatal period, when recurrent connectivity is strengthening. Selectively activating or silencing of these neurons in neonates enhanced or suppressed recurrent synaptic strength, respectively. Thus, embryonically TRAPed piriform neurons represent an interconnected hub-like population whose activity promotes recurrent connectivity in early development.

## INTRODUCTION

During development, neuronal activity emerges before neural circuits are mature. At the circuit level, such activity is known to play a key role in refining feedforward synaptic connections that propagate sensory information into the brain^1–6^. As a specific example, in the developing vertebrate visual system, spontaneous retinal waves prior to vision onset are required for the segregation of eye-specific retinal input in target regions^2^ and interact with axon guidance molecules to establish precise topographic map^7,8^. Downstream of these afferent inputs, diversity in the recurrent circuit architectures underlies the vast catalog of transformations performed by CNS circuits such as gain modulation, sensory integration, and working memory^9–17^. For example, recurrent excitatory connectivity and feedback inhibition in the adult mouse olfactory cortex is necessary for stable representation of odor information^18–20^. Features of recurrent connectivity in olfactory cortex also establish a discrete ensemble representation of different odors and shape temporal dynamics that provide rich information about incoming sensory activity^18,21–23^.

While neural activity is known to regulate development of individual neurons such as cell type differentiation, apoptosis, and morphological maturation^24–27^, how such activity shapes the development of network features such as recurrent connectivity has not been directly explored. Indeed, instances of spontaneous network activity have been described in some of developing recurrent circuits such as the neonatal hippocampus and temporal cortices^28–31^. However, such studies do not explicitly explore how spontaneous activity affects the development of connectivity within these networks. Because recording neuronal populations in developing animals has historically been challenging, it is also an open question whether specific cell populations regulate activity in such networks *in vivo*^32,33^. Lastly, it is unclear if activity in developing recurrent circuits corresponds to transformation of sensory input or to sensory-independent developmental processes.

Neuronal activity in developing mammals has been difficult to study due both to technical limitations (e.g., inaccessibility of developing nervous system *in utero* and instability of neonatal crania for head-fixed recordings) and biological constraints (e.g., low spiking activity of developing neurons^34^). Here, we leveraged the TRAP method^35,36^ to perform the first brain-wide survey of neuronal activity in the developing mouse brain, focusing on embryonic stages where activity has only been observed in a select group of more readily accessible brain regions^6,37–39^. TRAP enabled non-invasive and permanent labeling of embryonically active neurons, allowing us to investigate their role in network development over multiple subsequent stages. Following embryonic TRAP, we performed tissue clearing and whole-brain imaging, and identified the primary olfactory (piriform) cortex as one of the most abundantly labeled brain regions.

Olfaction is essential to neonatal survival^40–43^, and the piriform cortex depends heavily on its recurrent circuit architecture to maintain ensemble representation of odors^18–20,22,23,44,45^. Neuronal activity in the embryonic piriform cortex could therefore be important for both neonatal ethology and recurrent circuit development. Using a combination of *ex vivo* and *in vivo* electrophysiological approaches, we find that embryonically TRAPed piriform neurons are a preferentially interconnected hub-like population that exhibits broad connectivity with the rest of the piriform network. During neonatal stages, peaks in activity of these neurons lead synchronized population activity. Chronic *in vivo* activation of this population in neonates promotes persistent increases in recurrent connectivity, while silencing this population reduces recurrent connectivity. Together, these findings suggest that embryonic activity in the piriform cortex contains a hub-like neuronal population whose activity helps establish recurrent circuit architecture in the developing mouse brain.

## RESULTS

### A TRAP2-based whole-brain screen of active neurons during late embryogenesis

To comprehensively identify brain regions enriched for active neurons during development, we applied the TRAP2 system^36,46^ to embryonic mouse brains (**Figure 1A**). We administered the activating drug 4-hydroxytamoxifen (4OHT) at embryonic day 18 (E18) to pregnant dams double transgenic for *TRAP2* and *Ai14* (a tdTomato Cre reporter^47^) to permanently label E18-active neurons (**Figure 1B**). After TRAPed embryos grew to adulthood (>P40), we used the whole-brain clearing method iDISCO^48^ and cell quantification pipeline ClearMAP^48^ to identify E18-TRAPed brain regions. We found many regions with substantial numbers of labeled neurons, including basal ganglia, amygdala, thalamus, hypothalamus, and brainstem (**Figure 1C, D; Table S1**). Several of these regions have been the focus of studies of early neuronal activity, such as striatal neurons in the postnatal development of corticostriatal connectivity^49^, subplate neurons in developing cortical microcircuits^50^, and fetal suprachiasmatic nucleus in response to maternal circadian rhythms^51^. Several dorsal cortical regions that are known to have active neurons in early development were also labeled. In addition, we identified transient neuronal populations such as Cajal-Retzius neurons by also collecting E18-TRAPed brains in early postnatal stages (P4) (**Figure S1A**)^50,52^. Overall, embryonic neuronal activity in most of our identified regions had not been previously reported and could be subjects of future neurodevelopmental studies.

**Figure 1.**
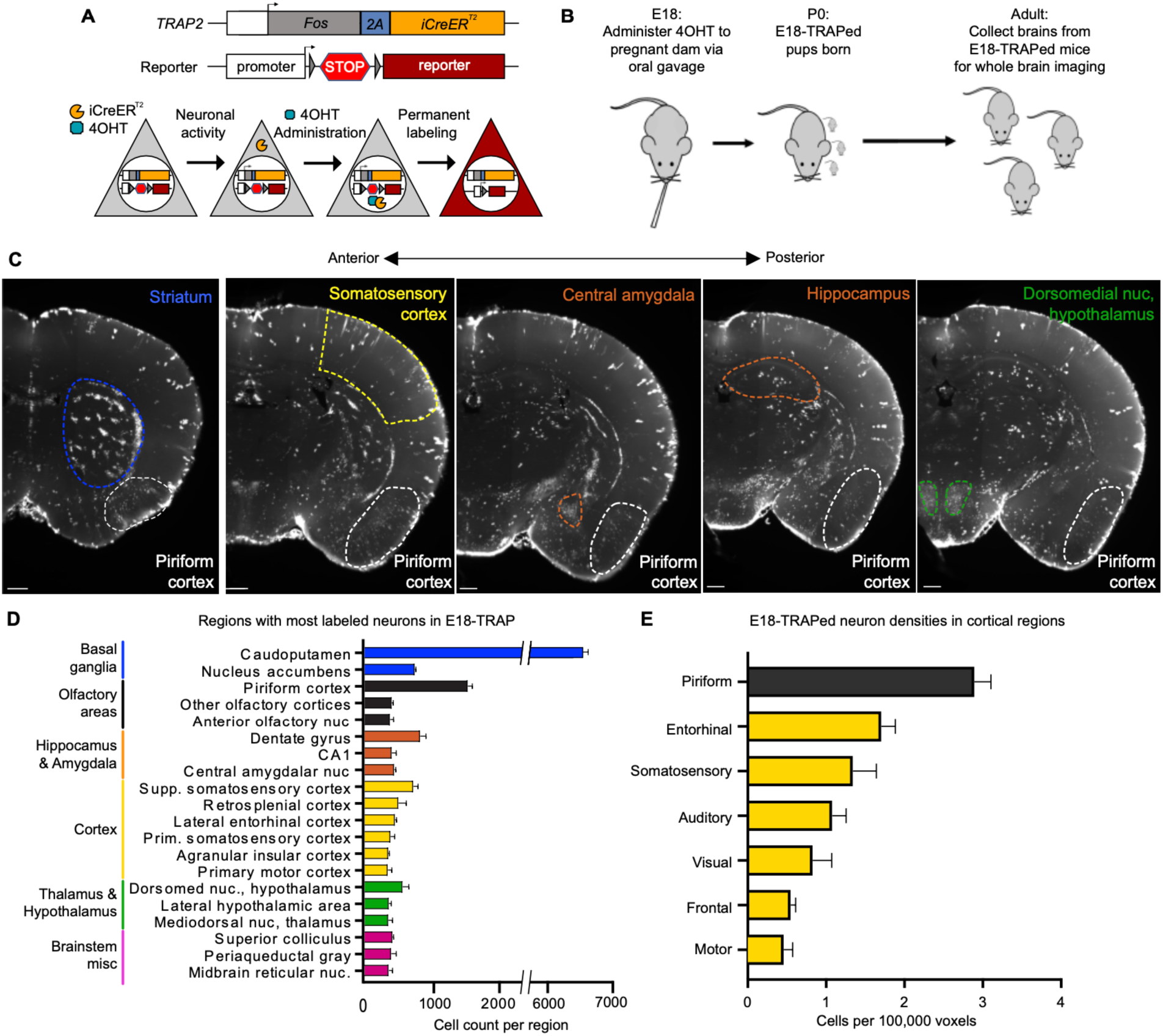
E18-TRAP captures active neurons in diverse brain regions during prenatal development. (A) Schematic of the TRAP strategy and the *TRAP2* allele^36^. Neuronal activity and 4-hydroxytamoxifen (4OHT) application induce transcription and nuclear translocation of CreER, respectively, causing recombination that removes the stop between two *loxP* sites (triangles) and leading to permanent expression of the reporter. Arrow, transcriptional start. (B) Schematic of TRAPing embryonic neuronal activity via oral gavage of 4OHT to pregnant dam. (C) Example images along anterior-posterior axis of E18-TRAPed brain cleared and imaged at P40. Piriform cortex and other abundantly TRAPed regions are circled. Note that most brightly labeled large cells (e.g., in neocortex and hippocampus) are astrocytes. Scale bar, 100 μm. (D) Average cell counts of the top 20 E18-TRAPed brain regions (n = 3 mice). Error bars, SEM. (E) Average cell densities of the cortical regions in E18-TRAPed brain (n = 3 mice). Error bars, SEM. See additional data in Figure S1 and Table S1.

The piriform cortex contained the greatest density of E18-TRAPed neurons among cortical regions (**Figure 1E**), and the second-most abundantly labeled regions brain-wide (**Figure 1C, D; Table S1**). TRAP at earlier developmental stages demonstrated that piriform cortical activity was present as early as E14, although the number of TRAPed neurons was substantially higher when active neurons were captured at E18 (**Figure S1B, C**). Several other olfactory related regions were also among the regions that contain most E18-TRAPed cells (**Figure 1D, Table S1**).

Previous *ex vivo* studies in perinatal and early postnatal cortical brain slices observed that temporal cortical regions containing piriform cortex were more spontaneously active than other cortical regions in early development^38,53^, and fiber photometry in neonates identified active temporal cortical regions with cortex-wide calcium waves^28,30^. Because of the ethologically important role of neonatal olfaction^40–43^ and role of recurrent intracortical connectivity in adult piriform^18,19,45^, we focus the rest of this study on investigating the properties and function of E18-TRAPed piriform neurons.

### Basic properties and potential roles of E18-TRAPed piriform neurons in a developing recurrent network

To validate that embryonic Fos-expressing neurons (the basis of TRAP) are electrically active, we performed RNAscope on E18 embryonic piriform cortex of wild-type mice with multiple immediate early genes, including *Fos*, *Arc*, and *Npas4*, that have different activation pathways downstream of electrical activity^54^. We found co-expression of multiple immediate early genes in most piriform neurons (**Figure 2A, B**). This suggests that labeling of these neurons is likely due to activity rather than activity-independent pathways, especially given that *Npas4* expression is highly specific for neuronal activity^55^. To characterize the distribution of excitatory vs. inhibitory neurons among the E18-TRAPed population, we performed RNAscope on E18-TRAPed neurons at P3.5 with a *Vgat* probe (encoding vesicular GABA transporter, a GABAergic neuronal marker). We found that E18-TRAPed neurons had similar proportions of *Vgat*+ neurons as the general piriform cortex population (∼15%; **Figure 2C, D**).

**Figure 2.**
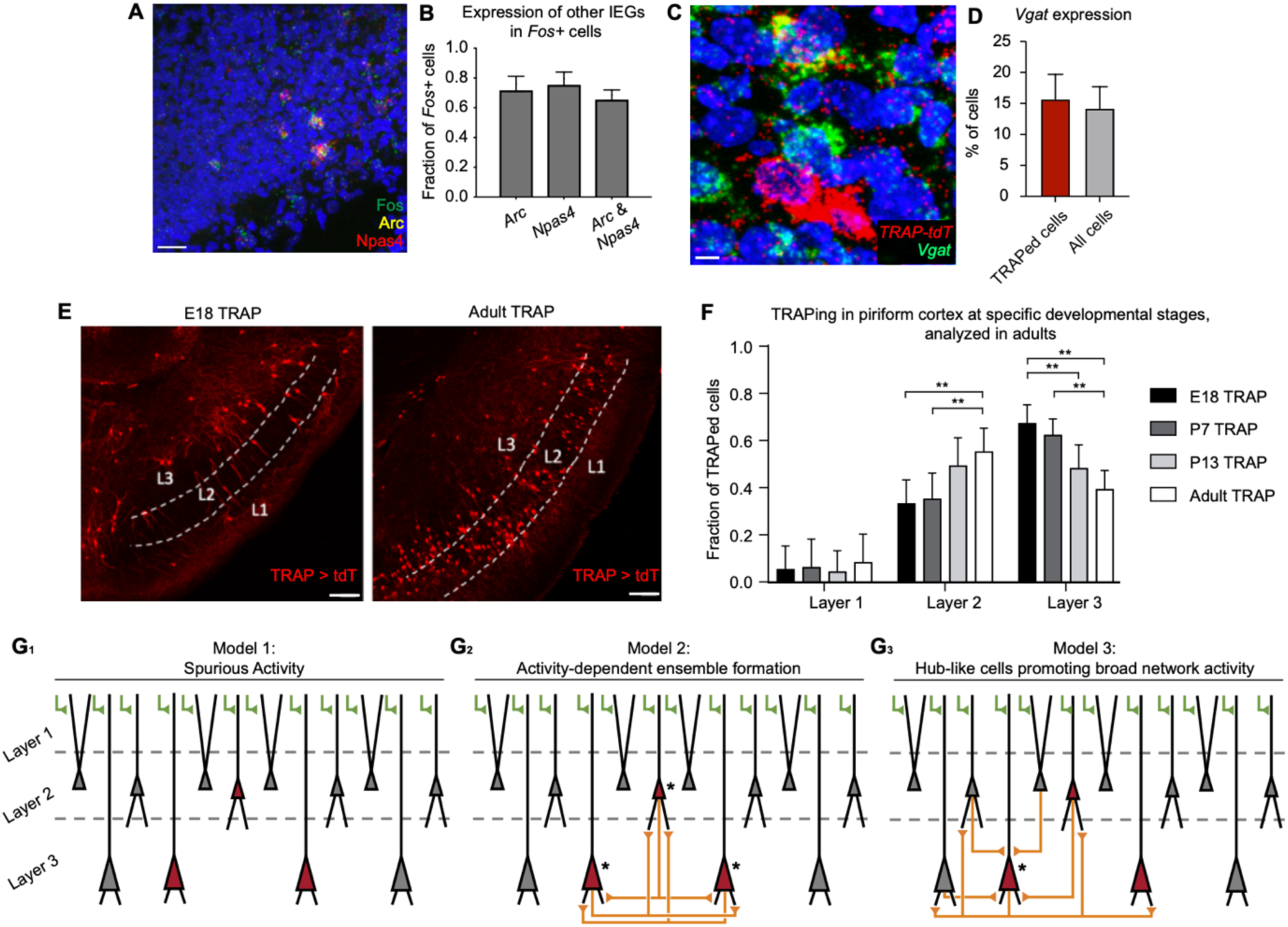
Characterization and potential developmental roles of E18-TRAPed neurons in piriform cortex. (A, B) Example (A) and quantification (B) of RNAscope for co-expression of immediate early genes *Arc* and *Npas4* in *Fos*-expressing cells in E18 piriform (n = 4 mice). Error bars, SEM. Scale bar, 20 μm (C, D) Example (C) and quantification (D) of RNAscope for expression of *Vgat* in E18-TRAPed cells, collected in P3.5 brains (n = 6). Error bars, SEM. Scale bar, 4 μm (E, F) Example (E) and distribution (F) of active neurons across layers of piriform cortex when TRAPed at different ages (adult indicates >P40). Quantification of cells in animals TRAPed at different ages were all performed in adult mice (n = 4 mice per age; **p<0.005, 2-way ANOVA with post-hoc Tukey HSD). Error bars, SEM. Scale bar, 100 μm. (G) Three models for potential roles of embryonically TRAPed neurons in piriform network development. See text for more details. Asterisks indicate neurons of particular interest within each model. See additional data in Figures S1.

Histological analysis revealed that E18-TRAPed piriform populations were enriched in layer 3 of the piriform cortex compared with adult-TRAPed piriform populations (**Figure 2E, F, S1F**)^22^. TRAP in postnatal stages (e.g., P7 for an early postnatal stage, and P13 for a later postnatal stage after eye-opening and increased environmental exploration^56^) reveals similar layer distributions, with a gradual shift toward layer 2 dominance when TRAP was performed in adults (**Figure 2F**). Within layer 3, TRAPed populations comprise similar proportions of deep pyramidal and multipolar neurons (**Figure S1D, E**). Little is known about the function of either of these cell types during development or in the mature piriform cortex, with existing studies typically describing broader odor tuning, decreased olfactory bulb input, and increased recurrent connectivity compared to the more commonly studied layer-2 piriform neurons^20,22,57–60^. The overrepresentation of TRAPed cells in layer 3 in early developmental stages suggests that they may play a role in network development.

Lastly, most E18-TRAPed neurons were present and already labeled in piriform cortex by birth (**Figure S1G**), confirming that most of these neurons are indeed present and active within piriform before birth. The fraction of E18-TRAPed neurons in very deep layer-3 was smaller at birth than in adults (**Figure S1G**), suggesting that some of these neurons may have migrated to piriform postnatally and are less likely to be relevant to early activity in piriform. Thus, neurons at these depths were excluded from analysis in remaining experiments.

Because the piriform cortex has abundant recurrent connectivity and odors activate distributed ensembles of recurrently connected neurons in adults, we were interested in how early activity may shape this. We considered three possible scenarios for embryonically active neurons in the development of the piriform cortex network (**Figure 2G**). First, TRAPed neurons could simply represent a random population that happened to be active during the 4OHT application period; such spurious neuronal activity may not contribute to circuit development (**Figure 2G_1_**).

Second, embryonic activity may drive the self-assembly of active neuronal ensembles (**Figure 2G_2_**). Recurrent connectivity is thought to facilitate an ensemble representation of odors in adult piriform, but what processes shape the maturation of these recurrent synapses and establishment of ensemble connectivity are understudied^18,19,22,23^. Given the importance of olfaction in neonates^40,42^, embryonic piriform activity can help establish preferential connectivity between co-active neurons via Hebbian plasticity^61^. Further, such ensembles may co-tune towards similar odors during olfactory imprinting, and strongly connected ensembles may facilitate consistent reactivation of neurons representing ethologically relevant odors.

Third, E18-TRAPed piriform neurons may constitute a highly active and broadly connected hub-like population whose activity may drive network synaptogenesis broadly throughout early development (**Figure 2G_3_**). Hub neurons with the ability to modulate synchronous population events have been observed in brain slices of developing recurrent networks, such as neonatal hippocampus^28,30,33,62^. However, their existence and function *in vivo* are not well-understood, and the role of their activity in circuit maturation has not been explored.

We note that these scenarios are neither necessarily mutually exclusive (e.g., between the 2^nd^ and 3^rd^ scenarios) nor do they contain all possibilities. Nevertheless, they serve here as a framework for directing the following brain slice and *in vivo* experiments to narrow down the possibilities.

### Preferential connectivity of E18-TRAPed piriform neurons in *ex vivo* brain slices

Both the ensemble (**Figure 2G_2_**) and hub neuron (**Figure 2G_3_**) models predict distinct patterns of synaptic connectivity in E18-TRAPed versus nonTRAPed cells in the piriform cortex. To test this prediction, we performed a series of experiments in acute brain slices from P2–10 E18-TRAPed mice. For these experiments, we obtained whole-cell, voltage-clamp recordings from TRAPed and nonTRAPed neurons, holding cells at –70 mV to isolate excitatory postsynaptic currents (EPSCs).

In the first set of experiments (**Figure 3A–D**), we tested if E18-TRAPed neurons receive more or stronger synaptic inputs by recording miniature excitatory postsynaptic current (mEPSCs) from TRAPed vs nonTRAPed neurons in acute brain slices from *TRAP2/TRAP2;Ai14/Ai14* neonates (**Figure 3A**). We found both mEPSC amplitude and frequency were significantly increased in E18-TRAPed neurons compared to nonTRAPed neurons, suggesting that they receive stronger and overall more synaptic inputs, respectively (**Figure 3B–D**).

**Figure 3.**
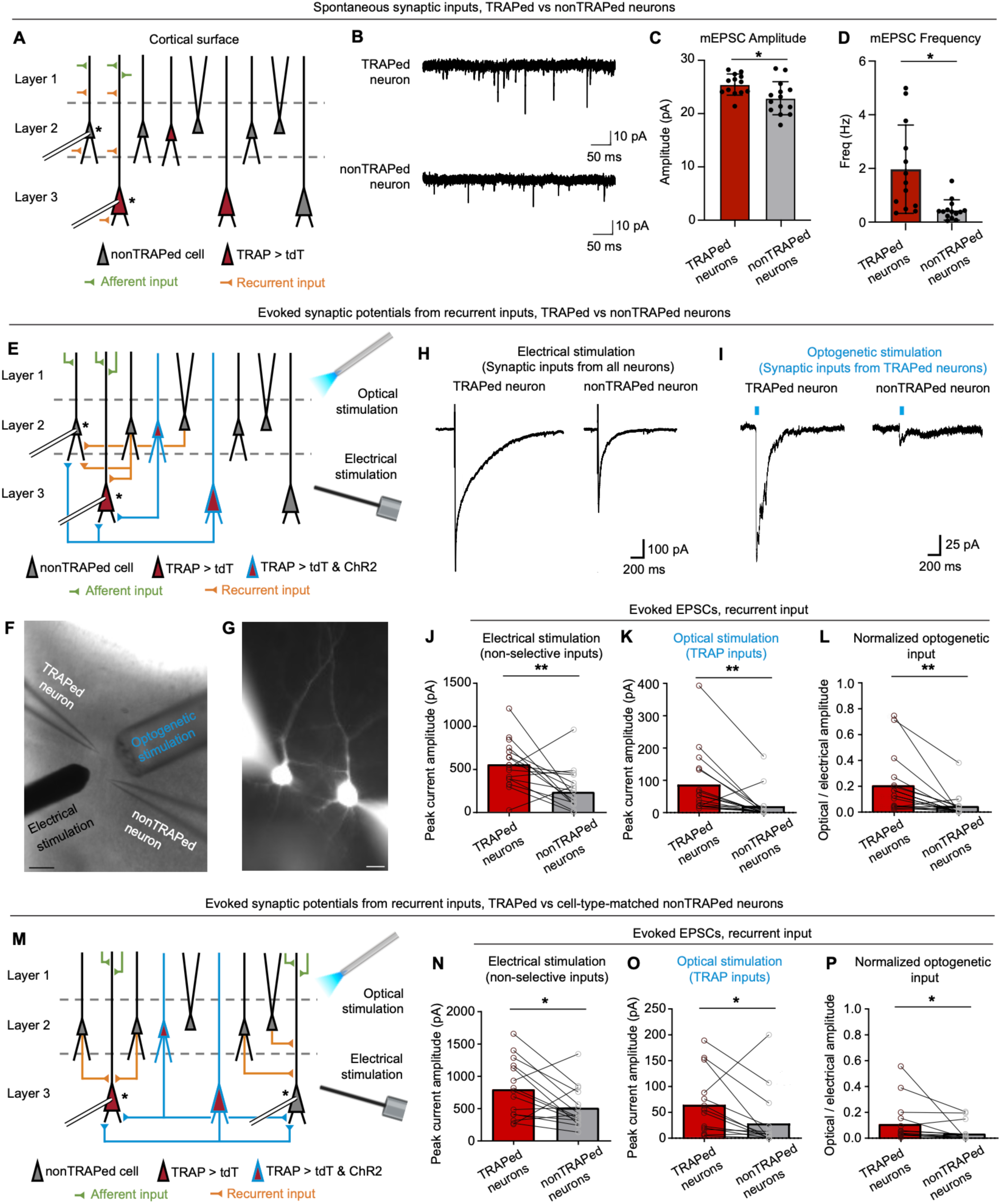
Synaptic connectivity of E18-TRAPed neurons in acute neonatal slice. (A) Schematic of investigating connectivity of E18-TRAPed neurons by recording mEPSCs. Patch-clamp recording was performed from TRAPed and nonTRAPed cells in the piriform cortex. Note that pyramidal neurons in layers 2 and 3 both receive olfactory bulb afferent input (green) at superficial layer 1, and recurrent input from other pyramidal neurons (orange) at deep layer 1 as well as layers 2 and 3 (B–D) Example mEPSC traces (B) and quantification of mEPSC amplitude (C) and frequency (D) of TRAPed and nonTRAPed cells in neonatal acute slice (n = 13 for TRAPed cells, n = 14 for nonTRAPed cells across 5 mice). Error bars, SEM. (E) Schematic of investigating connectivity of E18-TRAPed neurons with electrical and optogenetic stimulation. Asterisks indicate cells that are recorded. Recording from tdTomato (tdT)-positive but ChR2-negative TRAPed neurons in response to optogenetic stimulation allows recording of inputs from ChR2-positive TRAPed cells to such ChR2-negative TRAPed cells. (F) Representative image of electrical and optogenetic stimulation of simultaneously recorded E18-TRAPed and nonTRAPed cell in slice. Scale bar, 100 μm. (G) Example of a pair of TRAPed and nonTRAPed neurons in dual-patch setup, filled with Alexafluor 555 dye for morphological visualization. Scale bar, 15 μm. (H) Example averaged trace of 5 evoked postsynaptic currents in a TRAPed and a nonTRAPed cell, following electrical stimulation of all synaptic inputs. (I) Example averaged trace of 5 evoked postsynaptic currents in a TRAPed and a nonTRAPed cell following optogenetic stimulation of inputs from TRAPed neurons (J, K) Summary of peak current amplitude under electrical stimulation of recurrent inputs from all neurons (J) or optogenetic stimulation of recurrent inputs from TRAPed neurons (K). n = 18 pairs per condition across 5 mice. (L) Summary of optogenetic peak current (input from TRAPed neurons) normalized by electrically evoked peak current (input from all neurons) for recurrent input for cells in (J, K). (M) Schematic of comparing connectivity of E18-TRAPed neurons with cell-type-matched nonTRAPed cells with electrical and optogenetic stimulation. Asterisks indicate cells that are recorded. (N, O) Summary of peak current amplitude under electrical stimulation of recurrent inputs from all neurons (N) or optogenetic stimulation of recurrent inputs from TRAPed neurons (O) to TRAPed vs cell-type-matched nonTRAPed neurons. n = 17 pairs per condition across 12 mice. (P) Summary of optogenetic peak current (input from TRAPed neurons) normalized by electrically evoked peak current (input from all neurons) for recurrent input for cells in (N, O). *p< 0.05, **p< 0.005, ***p< 0.001, paired t-test. See additional data in Figures S2 and S3.

Next, we tested if E18-TRAPed neurons were preferentially interconnected. For these experiments, we used *TRAP2/TRAP2;Ai14/Ai32* mice that express Cre-dependent tdTomato and/or ChR2-eYFP^47^. The transient expression and nuclear translocation of CreER^T2^ in TRAP make Cre-dependent recombination somewhat stochastic. This resulted in subsets of TRAPed neurons that express only tdTomato, only ChR2, or both tdTomato and ChR2 (**Figure S2A, B**). We recorded from TRAPed neurons expressing only tdTomato to avoid evoking direct ChR2- mediated currents in the recorded cells (**Figure 3E–G**). For each recorded TRAPed neuron, we simultaneously recorded from a nonTRAPed piriform neuron within the same slice. To selectively stimulate synaptic input from TRAPed neurons, we used brief light pulses to activate the axons of TRAPed neurons expressing ChR2. We then measured the resulting EPSCs in simultaneous recordings from the TRAPed and nonTRAPed neurons. In addition, we used a stimulating electrode to activate a larger and unbiased population of inputs onto the recorded neurons (hereafter abbreviated as ‘non-selective input’). After recording baseline EPSCs, we applied the GABA_B_ receptor agonist baclofen to the extracellular solution to block all but afferent sensory inputs, as recurrent but not afferent axons express GABA_B_ receptor at their axon terminals^63,64^. This allowed us to distinguish between, and thus compare the strengths of both afferent and recurrent inputs to TRAPed and nonTRAPed cells.

E18-TRAPed piriform neurons exhibited larger postsynaptic currents than nonTRAPed cells in response to both electrical and optogenetic stimulation, as seen by the example traces (**Figure 3H, I**) and quantifications (**Figure 3J–L; Figure S2C, D**). Specifically, by comparing responses to electrical stimulation before and after baclofen application, we found that peak currents of both recurrent and afferent inputs were significantly higher in TRAPed neurons compared to control neurons (**Figure 3J; Figure S2C**). This suggests that E18-TRAPed neurons receive more non-selective recurrent synaptic input than nonTRAPed neurons, consistent with the mEPSC analysis (**Figure 3A–D**). The high variance in evoked synaptic responses is consistent with piriform cortical neurons having substantial differences in their afferent and recurrent input strength^64^. Following optogenetic stimulation of inputs from other TRAPed cells, peak currents of both recurrent and afferent inputs were also significantly higher in TRAPed neurons compared to control neurons (**Figure 3K; Figure S2D)**. These findings suggest that TRAPed populations within the piriform cortex are preferentially interconnected.

Could the stronger responses observed in TRAPed neurons simply reflect an overall increase in synaptic inputs onto these neurons? To test this, we normalized the optogenetically-evoked EPSCs from TRAPed neurons by the non-selective electrical evoked current. After normalizing, we found that TRAPed neurons still received significantly greater recurrent input from other TRAPed neurons (**Figure 3L**), validating that E18-TRAPed neurons are preferentially interconnected.

Because TRAPed cells were most often located in deep piriform cortical layers, one possibility is that the differences in synaptic properties we observed were not specific to TRAPed cells but rather to deep layer cell types. To control for this (**Figure S2G–I**), we performed a third set of experiments (**Figure 3M–P**) in which we simultaneously recorded from a tdTomato-positive TRAPed neuron and a cell-type-matched nonTRAPed neuron within the same layer, with similar morphological (as judged by dye fill, see **Figure 3G**) and electrophysiological properties (e.g., membrane capacitance, resistance) (**Figure 3M**). Even when controlling for cell type, we found that TRAPed neurons received significantly stronger recurrent input in response to electrical (**Figure 3N**) and optogenetic (**Figure 3O**) stimulation, compared to nonTRAPed neurons. Interestingly, afferent inputs were no longer significantly different under both electrical and optogenetic stimulations (**Figure S2E, F**), suggesting that while TRAPed cells uniquely receive increased recurrent inputs, the increased afferent input may be accounted for by layer or cell-type differences. Importantly, TRAPed neurons still received stronger normalized recurrent input (optogenetic over electrical stimulation) than nonTRAPed neurons (**Figure 3P**), indicating that preferential connectivity between TRAPed neurons occurs even comparing with neighboring cell-type matched control cells.

In a fourth set of mice, we increased the threshold for TRAPing active neurons (**Figure S3A**). TRAPed populations were now substantially sparser and likely represent the most active neurons expressing the highest levels of Fos. Sparser TRAPed populations resulted in many fewer ChR2-expressing axons. Consequently, optogenetic stimulation of TRAPed neurons failed to evoke postsynaptic responses in most control neurons. However, many TRAPed neurons still exhibited robust postsynaptic responses, whereas only one nonTRAPed neuron exhibited any response (**Figure S3B, C**). These data suggest that the increased connectivity between TRAPed neurons is robust to experimentally defined TRAPing thresholds.

Lastly, we grouped the responses of all recorded neurons to by age. We found the strength of recurrent synaptic responses evoked by electrical stimulation increased significantly over the first two postnatal weeks (**Figure S2J**). However, there was no change in the strength of optogenetically-evoked synaptic responses across the same period (**Figure S2K**). This suggests that while general recurrent connectivity strengthens during this early postnatal period, output from TRAPed neurons to other piriform cells may be mostly saturated by this time. This positions the TRAPed population to play a key role in network maturation during early postnatal development.

Together, these findings suggest that E18-TRAPed piriform cortex neurons receive increased levels of recurrent inputs within piriform and exhibit disproportionately higher interconnectivity with other TRAPed neurons. TRAPed cell connectivity is also established early, whereas broad recurrent connectivity within piriform continues to strengthen postnatally. These findings suggest that E18-TRAPed cells have connectivity patterns consistent with both interconnected ensembles (**Figure 2G_2_**) and broadly connected hub-like populations (**Figure 2G_3_**), but argue strongly against TRAP simply capturing spuriously active neurons (**Figure 2G_1_**).

### Single-unit recordings *in vivo* revealed broad odor tuning of E18-TRAPed piriform neurons

Differentiating between the ensemble and hub neuron model requires observation of neuronal dynamics of E18-TRAPed neurons in response to odors and spontaneous activity epochs, which must be measured *in vivo*. Therefore, we established Neuropixels recordings in awake head-fixed P6–10 mice, which enabled simultaneous recordings of approximately 50–150 piriform neurons (**Figure 4A, S4A**). By performing these recordings in neonates, we aimed to capture potentially transient odor-tuning (e.g., to maternal odors), developmentally unique features of TRAPed neurons, or general dynamics of developing cortical networks (e.g., influenced by GABA acting as an excitatory neurotransmitter^65^) that may be lost in recordings on adult piriform.

**Figure 4.**
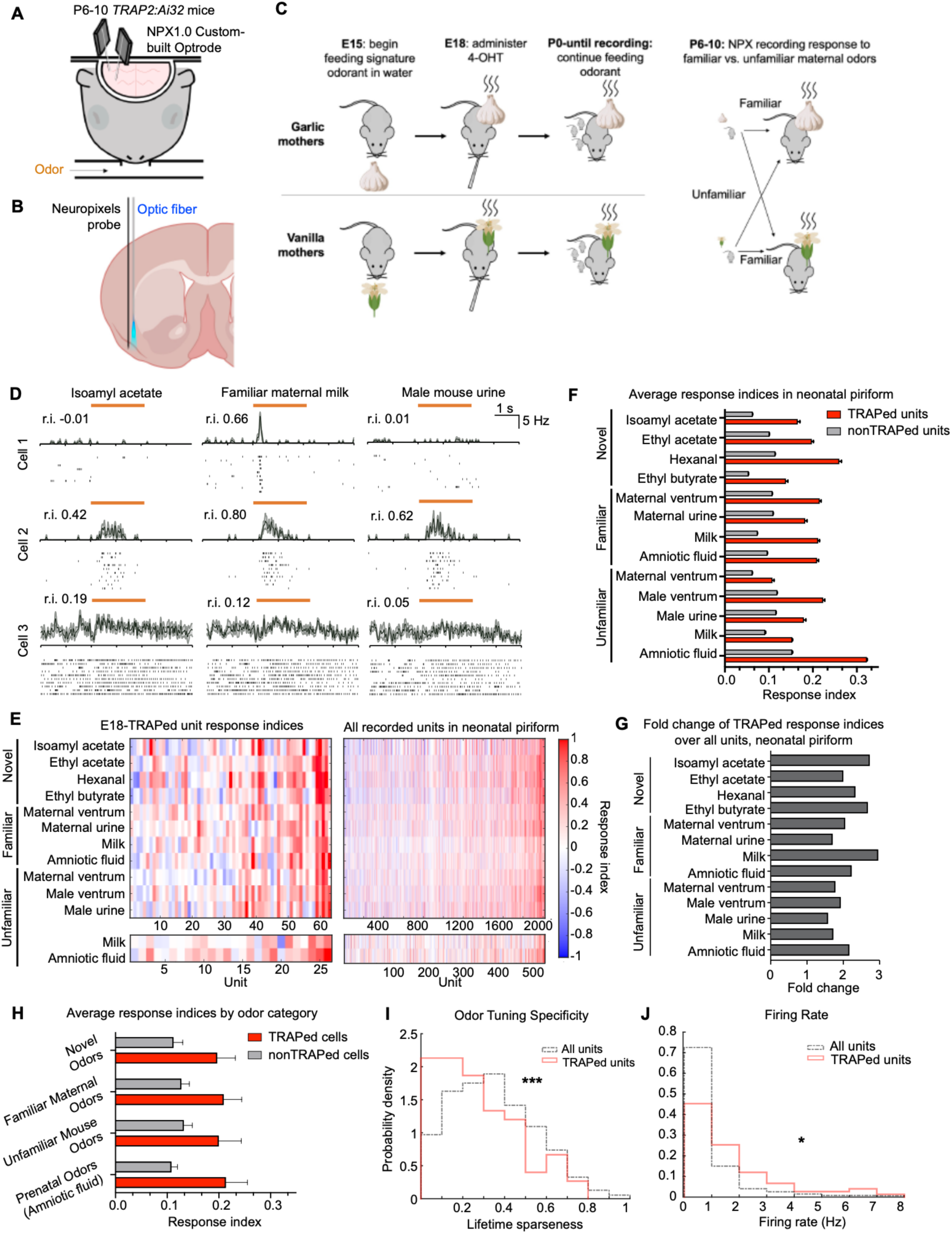
*In vivo* odor-response characteristics of E18-TRAPed neurons in neonatal piriform cortex. (A) Schematic of dual Neuropixels optrode recording in piriform cortex of awake P6–10 *TRAP2/TRAP2;Ai32/Ai32* mice. (B) Schematic of custom-built Neuropixels optrodes for recording and optotagging E18-TRAPed cells. (C) Experimental design to rear E18-TRAPed neonates in distinct odorant environments created by feeding signature odorants to dams in perinatal stages. (D) Example odor-evoked average peristimulus time histograms (PSTHs) and spike trains for 10 odor trials in 3 neonatal piriform units. Response index (r.i.) is displayed for each cell–odor pair. Orange bar indicates odor presentation epoch. (E) Response indices of individual E18-TRAPed units and all recorded units in neonatal piriform to 10 presentations of each odorant. A subset of recordings included Unfamiliar Milk and Amniotic Fluid, and are displayed separately. (F) Average response indices of E18-TRAPed vs all units recorded in neonatal piriform. Error bars, SEM across units. (G) Fold change of response indices of TRAPed compared to nonTRAPed units in neonatal piriform. (H) Average response indices across categories of odors of E18-TRAPed vs nonTRAPed units recorded in neonatal piriform. Error bars, SEM across units (I, J) Lifetime sparseness (I; See Methods) and baseline firing rate (J) of TRAPed vs nonTRAPed units in neonatal piriform. For panels E–J, n_TRAP_ = 63 units, n_Total_ = 2020 units, across 20 neonatal mice, except for Unfamiliar Milk and Unfamiliar Amniotic Fluid, where n_TRAP_ = 26 units, n_Total_ = 564 units (these units are a subset of the 63 TRAPed and 2020 total units). See additional data in Figures S4 and S5.

We used *TRAP2/TRAP2;Ai32/Ai32* animals to “optotag” E18-TRAPed piriform neurons (**Figure 4B; Figure S4B–D**). We recorded a total of 2020 neurons in 20 neonates, 63 of which were E18-TRAPed neurons identified by optotagging. As a comparison, we also recorded 864 neurons in 4 adult mice, 17 of which were E18-TRAPed neurons (**Figure S4G–K**). Although TRAPed neurons were sparse (average 4 per mouse), we were able to examine co-tuning to specific odorants and general physiological features of individual TRAPed neurons. We applied novel odorants (pure chemicals) as well as ethologically relevant odors from familiar and unfamiliar dams fed with signature odorants during perinatal stages (**Figure 4C**). Neurons recorded in neonatal piriform demonstrated diverse odor response profiles (**Figure 4D**). To quantify a unit’s firing to a given odor, we calculated the response index (r.i.; defined as 2 x auROC – 1, where auROC stands for area under the receiver operating characteristic curve) of a cell– odor pair from the response during a 2-second (2s) odor presentation compared to a 2s baseline before odor onset, with a response index of +1 corresponding to consistent excitation and –1 to suppression^21^.

E18-TRAPed piriform populations had consistently higher response indices across all odorants compared to nonTRAPed neurons (**Figure 4E–H**). Individual E18-TRAPed neurons also had broader tuning specificity as measured by lifetime sparseness (**Figure 4I**), a metric of odor response specificity where 0 indicates complete non-specificity and 1 indicates maximal specificity towards one odor. In addition to their broad tuning, E18-TRAPed neurons also had higher baseline firing rates (**Figure 4J**) compared to nonTRAPed neurons. However, E18-TRAPed piriform neurons did not appear to be preferentially co-tuned towards any specific odors or odor sets (**Figure 4E–G**). This was true for both odor sets from familiar mouse mothers whose odors may have facilitated olfactory imprinting, odors from unfamiliar mice, prenatal odors (amniotic fluid), and a control set of novel odors (**Figure 4F–H**). Given previous evidence for prenatal olfactory experience^42,66,67^, a subset of experiments also included amniotic fluid to examine if the E18-TRAPed piriform neurons might have responded to their specific embryonic olfactory environment. However, TRAPed neurons were not preferentially responsive to these odors (**Figure 4F–H**). TRAPed neurons also did not respond earlier or with more prolonged activity in response to odors compared to nonTRAPed neurons, whether measured in neonates or in adults (**Figure S4E–H**). Lastly, we found that near-threshold optogenetic stimulations of E18-TRAPed neurons, while mildly increasing the baseline firing rate of nonTRAPed cells, did not increase the odor response or alter odor response kinetics of nonTRAPed cells (**Figure S5**), suggesting that E18-TRAPed neurons do not directly amplify odor responses at early postnatal stage.

Interestingly, in the adult piriform cortex (**Figure S4I, J**), differences in both the lifetime sparseness (or tuning) and baseline firing rates between TRAPed and nonTRAPed neurons were no longer statistically significant. Odor responses in adult piriform units also skewed towards suppression compared to neonatal piriform units, consistent with maturation of inhibition after the early postnatal period (**Figure S4K**).

Because our recordings were limited to P6–10 mice (due to skull instability and low ChR2 expression for optotagging in pups younger than P6), it is possible that we missed an earlier role for E18-TRAPed neurons in neonatal olfaction. In a separate set of experiments, we used immediate early gene expression to probe responses to a familiar maternal versus novel control odor in younger mice. Using RNAscope for the activity-dependent genes *Fos* and *Arc*, we found that E18-TRAPed neurons did not respond differently to presentation of maternal versus control odors immediately after birth, at P3, or at P10 (**Figure S4L–N**). However, TRAPed neurons were more likely to express *Fos* or *Arc* across all conditions at birth and at P3 (**Figure S4O**), consistent with their broader tuning and higher baseline firing rate than nonTRAPed neurons at early postnatal stages (**Figure 4I, J**).

Together, these findings suggest that E18-TRAPed neurons do not represent ethologically relevant odors for neonates. While this does not rule out the possibility that some TRAPed neurons may be interconnected ensembles that co-tune to other odors, the broad responsivity and elevated baseline activity of E18-TRAPed neurons are more consistent with a central hub-like population in neonatal piriform cortex (**Figure 2G_3_**). Moreover, the absence of these characteristics in adulthood and the failure to enhance odor responses with TRAPed cell stimulation suggest that they may play a developmental function unrelated to the propagation of odor responses.

### Population activity analysis revealed hub-like properties for E18-TRAPed piriform neurons

If TRAPed neurons indeed behave as a hub-like population, we expect them to have increased connectivity with other piriform neurons *in vivo* and potentially play a role in organizing network activity^33^. To investigate functional connectivity *in vivo*, we took advantage of simultaneous recordings of large piriform populations and examined the spike timing correlation between all neurons within a single recording session to measure functional connectivity (**Figure 5A; Figure S6A**). We found that E18-TRAPed piriform neurons were more likely to form excitatory functional connections with other neurons (**Figure 5B_1_**). E18-TRAPed cells both had more leading functional connections (a proxy for sending synaptic outputs to other neurons) as well as more lagging functional connections (a proxy for receiving synaptic inputs from other neurons) (**Figure S6A, B**). Again, this preferential connectivity of E18-TRAPed neurons disappeared in adult animals (**Figure 5B_2_, S6C**). This is consistent with our *ex vivo* findings that TRAPed neurons receive stronger synaptic input from all neurons and demonstrates that TRAPed neurons have hub-like properties *in vivo* during neonatal stages.

**Figure 5.**
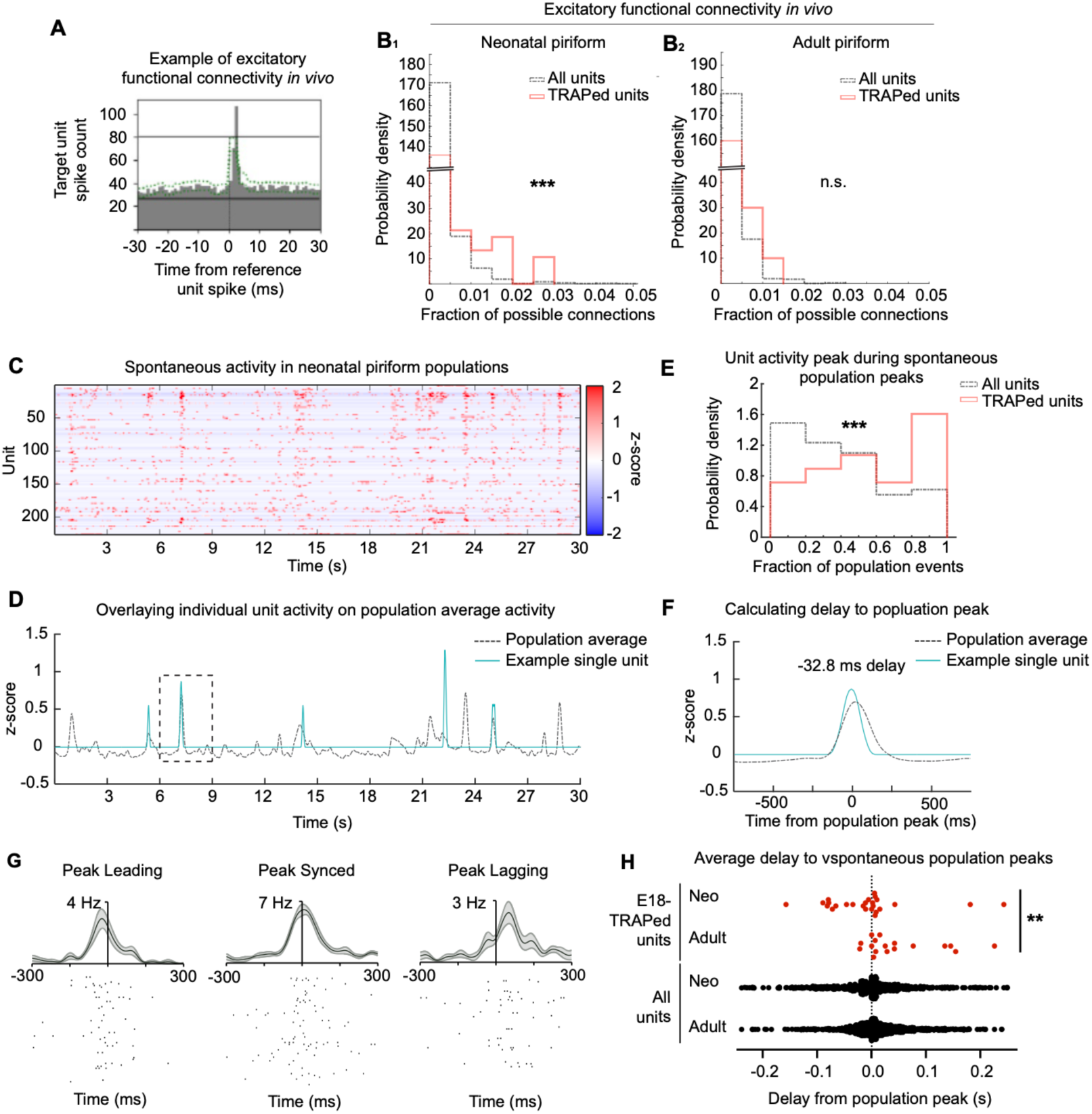
E18-TRAPed piriform neurons exhibit hub-like properties in neonatal piriform cortex. (A) Example cross-correlogram exhibiting excitatory functional connectivity. Dotted green trace represents significance via jitter method^79^.. Solid horizontal black lines within the plot indicate the upper and lower boundaries of significance. (B) Distribution of occurrence of excitatory functional connectivity in TRAPed vs all units in neonatal (B_1_) or adult (B_2_) piriform cortex measured as fraction of possible connections. Fraction of possible connections indicates the proportion of all recorded piriform cortical units within a mouse with which an individual unit had a functional connection, such that right-shifted distributions indicate that the population has increased functional connectivity. n_TRAP,Neonate_ = 63 units, n_Total,Neonate_ = 2020 units, n_TRAP_ = 17 units and n_Total_ = 864 units for adult mice; ***p<0.001 n.s., not significant, t-test with bootstrapping. (C) Example of individual unit activity over 30 s of spontaneous activity recording in one neonatal animal (colormap represents z-scored firing rates within 10-ms bins). (D) Average population z-score corresponding to epoch depicted in (C) overlaid with smoothed (over 200 ms) z-scored firing rate from a single unit. (E) Likelihood of E18-TRAPed vs all units having an activity peak during spontaneous population activity peaks, measured as fraction of population events. Fraction of population events indicates what proportion of all synchronized population activity peaks during which an individual unit also had an activity peak. n_TRAP_ = 26 units, n_Total_ = 564; ***p<0.0005, t-test with bootstrapping. (F) View of boxed region in (D) at higher temporal resolution demonstrating the delay between an individual unit activity peak (cyan trace) and population activity peak (black dotted trace). (G) Example PSTHs of three units centered around the peak of synchronized population events (vertical lines). (H) Distributions of average delay between peak of individual unit activity and peak of synchronized population events. n_TRAP,neonate_ = 26 units, n_TRAP,adult_ = 17 units, n_All,Neo_ = 564 units, n_All,Adult_ = 864 units across 5 neonatal mice and 4 adult mice; **p<0.005, unpaired t-test. Neo, neonate. See additional data in Figures S6 and S7.

Hub neurons in other developing networks have been shown to modulate network activity^32,33,68^. In our recordings of spontaneous activity in P6–10 neonates, we observed frequent occurrences of synchronized population activity (**Figure 5C, D**). E18-TRAPed neurons had significantly increased participation in these synchronized population events compared to nonTRAPed neurons (**Figure 5E**). Furthermore, the peaks of activity in individual E18-TRAPed piriform neurons tended to briefly precede the peaks in population activity (**Figure 5F–H**). This observation suggests a causal role for TRAPed neurons in evoking these bursts of population activity. In addition, this temporal offset argues against the increase in TRAPed neuron spiking during population bursts being simply due to their higher firing rates. Again, this phenomenon disappeared in the adult piriform (**Figure 5H, S6D**), suggesting that these neurons may play a role in initiating such events in the developing piriform network, perhaps by increasing overall excitatory drive within the network. This finding was unlikely to be caused by spurious sensory-driven responses, as TRAPed neurons do not fire earlier in response to odor presentation compared to other piriform cortical neurons (**Figure S4E**). The fact that TRAPed cells led spontaneous population activity but did not have shorter latencies to odor responses again suggests they play a role unrelated to processing of odor information.

Although our TRAPed neurons were distributed across all piriform layers, TRAPed neurons whose activity peaked before population events were predominantly located in layer 3, where many of the large and excitatory deep pyramidal neurons reside (**Figure S6E, F**). This is consistent with our observation that E18 and early postnatal TRAP showed that layer 3 neurons are preferentially active in early development.

Lastly, unsupervised clustering of piriform cortex units using *in vivo* anatomical and physiological properties demonstrated that E18-TRAPed cells in neonatal stages preferentially arise from three cell clusters, two of which are particularly sparse (**Figure S7A–D**; clusters 1 and 2). Both clusters exhibited properties consistent with hub-like cells, including increased firing rate, excitatory functional connectivity, and participation in synchronous population events (**Figure S7E–G**). Interestingly, while both clusters have broad odor tuning, cluster 1 units more notably led synchronous population events without having stronger odor-evoked responses, consistent with a sensory-independent hub-like population (**Figure S7H–J**). These findings reflect that while E18-TRAPed neurons are heterogenous, they are enriched for small subsets of cells with hub-like properties.

Together, these findings suggest that embryonically TRAPed neurons indeed have hub-like properties (**Figure 2G_3_**). In addition, the disappearance of increased functional connectivity, participation in population events, and leading synchronizing population peaks in adult piriform suggest that E18-TRAPed neurons may play a transient role in modulating network dynamics during early development.

### Silencing E18-TRAPed neurons reduces recurrent connectivity in neonatal piriform cortex

The concept of hub neurons has been discussed in several neonatal networks, with evidence for such neurons being reported both *in vivo* in neonates and *ex vivo* in neonatal slice^32,33^. However, how hub neurons affect the development of neuronal networks has not been explored. In our *ex vivo* physiology experiments, we found that the strength of recurrent connectivity in both TRAPed and nonTRAPed piriform neurons increased during the first postnatal week (**Figure S2J**). Given that the hub-like properties of E18-TRAPed neurons persisted into early postnatal stages, we asked if activity of E18-TRAPed neurons at these stages is required for the observed increase in recurrent connectivity. To do this, we expressed tetanus toxin (TeNT) in E18-TRAPed neurons to block their synaptic output (**Figure S8A**). We then obtained whole-cell recordings from randomly selected piriform layer 2 principal neurons in brain slices isolated from these mice, and measured EPSCs evoked by electrical stimulation (**Figure 6A, B**). We further isolated recurrent and afferent inputs by comparing recordings before and after adding baclofen, as previously described (**Figure 3**).

**Figure 6.**
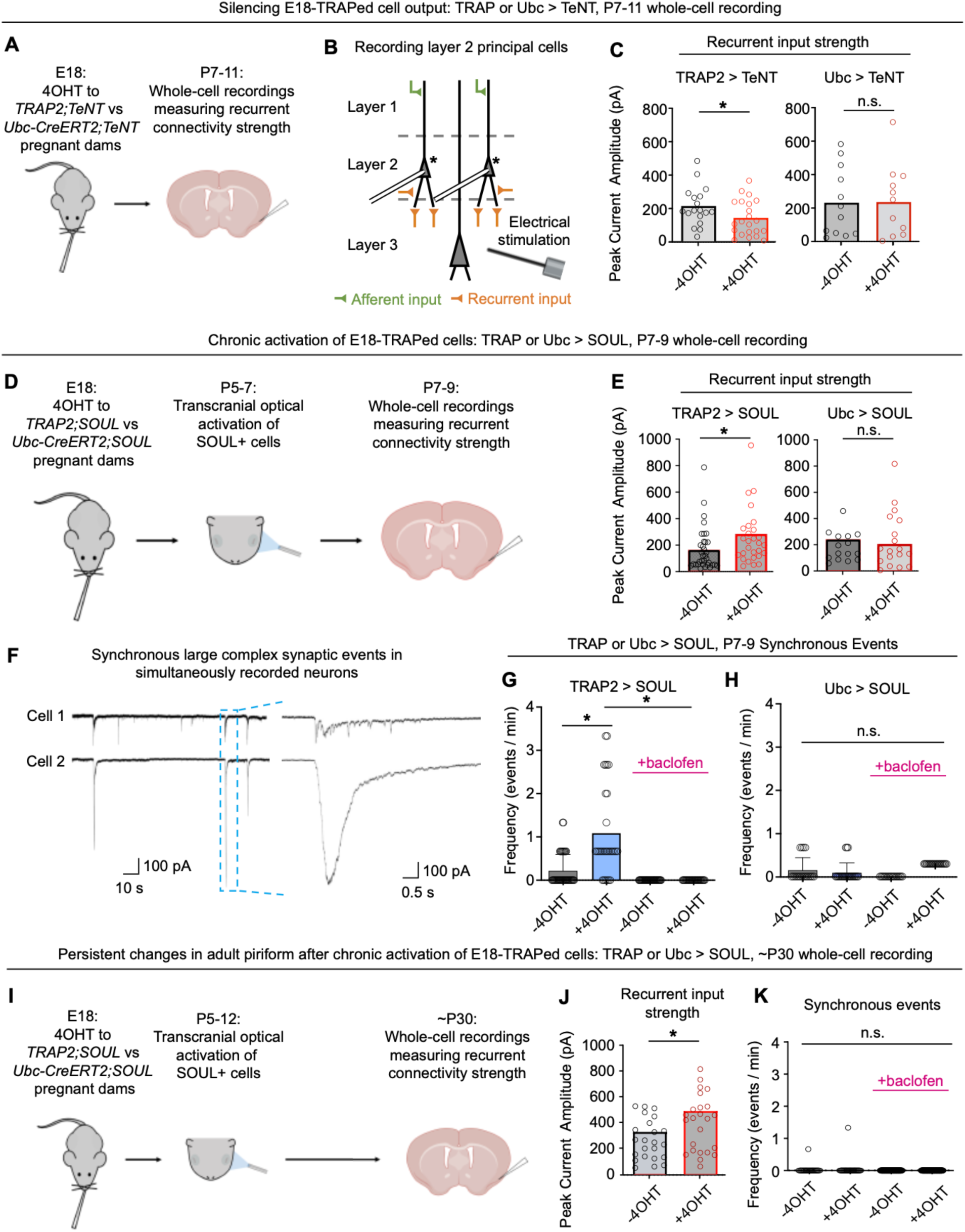
Bidirectional modulation of E18-TRAPed cell activity and resultant changes in recurrent connectivity in neonatal and adult piriform cortex. (A) Schematic of strategy of silencing E18-TRAPed cells in neonates. (B) Schematic demonstrating whole-cell recording of principal cells in piriform cortex. Cells with asterisks are recorded. (C) Plot of peak current amplitude of recurrent inputs in *TRAP2;LSL-TeNT* (TRAP > TeNT; *LSL*, *loxP-stop-loxP*) or *Ubc-CreER^T2^;LSL-TeNT* (Ubc > TeNT) animals compared to fostered littermates that did not receive 4OHT. n_-4OHT,TRAP_ = 22 cells across 5 mice, n_+4OHT,TRAP_ = 23 cells across 6 mice, n_-4OHT,Ubc_ = 11 cells across 3 mice, n_+4OHT,Ubc_ = 12 cells across 3 mice; *p<0.05, n.s. not significant; unpaired t-test. (D) Schematic of strategy of chronic activation of TRAPed cells in neonates. (E) Plot of peak current amplitude of recurrent inputs in stimulated *TRAP2;SOUL* (TRAP > SOUL) (L_1_) or *Ubc-CreER^T2^;SOUL* (Ubc > SOUL) (L_2_) animals compared to fostered littermates that did not receive 4OHT. n_-4OHT,TRAP_ = 25 cells across 5 mice, n_+4OHT,TRAP_ = 28 cells across 3 mice, n_-4OHT,Ubc_ = 13 cells across 2 mice, n_+4OHT,Ubc_ = 17 cells across 3 mice; *p<0.05, n.s. not significant; unpaired t-test (F–H) Example (F) and quantification (G, H) of frequency of synchronous large synaptic events in two simultaneously recorded neurons in neonatal piriform cortex following SOUL stimulation in neonatal *TRAP2;SOUL* mice (I; n_-4OHT,TRAP_ = 40 cells, n_+4OHT,TRAP_ = 30 cells) or *Ubc- CreER^T2^;SOUL* mice (J; n_-4OHT,Ubc_ = 18 cells, n_+4OHT,Ubc_ = 24 cells). *p<0.05, n.s. not significant; one-way ANOVA with post-hoc Tukey HSD. (I) Schematic of strategy of chronic activation of TRAPed cells in neonates followed by recording in adult piriform cortex (J, K) Plot of peak current amplitude of recurrent inputs (J) in stimulated *TRAP2;SOUL* (TRAP > SOUL) animals compared to fostered littermates that did not receive 4OHT. Quantification of frequency of synchronous large synaptic events (K) in two simultaneously recorded neurons in adult piriform cortex following SOUL stimulation in neonatal *TRAP2;SOUL* mice (M). n_-4OHT,TRAP_ = 23 cells across 4 mice, n_+4OHT,TRAP_ = 24 cells across 4 mice; *p<0.05, **p<0.005, n.s. not significant; unpaired t-test. See additional data in Figure S8.

TeNT expression in E18-TRAPed neurons led to a small but significant reduction of EPSC amplitude of recurrent but not afferent inputs to layer 2 principal neurons compared to control (TRAP animals without 4OHT injection to label active neurons) (**Figure 6C, S8G,** left panels). Given the very small number of E18-TRAPed neurons (∼1% of total piriform neurons; **Figure S1F**) compared to vast number of nonTRAPed neurons in the piriform cortex, it is unlikely that the observed reduction in EPSC amplitudes in randomly selected layer 2 principal cells was simply due to the loss of direct input from silenced E18-TRAPed neurons. However, to test this directly, and to ensure that the reduction in recurrent connectivity was specifically due to silencing TRAPed neurons, we performed parallel experiments using *Ubc-CreER^T2^*to express TeNT in similar number of random piriform cells (see **Figure S8D, E** described below). Now, TeNT did not affect peak EPSC amplitudes of recurrent or afferent inputs to layer 2 principal neurons (**Figure 6C, S8G**, right panels). Together, these data indicate that the activity of E18-TRAPed neurons is necessary for the maturation of recurrent connectivity in the piriform cortex, disruption of which reduces the strength of recurrent input across the piriform cortex.

### Activation of E18-TRAPed piriform neurons during neonates leads to persistent increase in piriform connectivity

To test if increasing the activity of E18-TRAPed neurons strengthens recurrent connectivity, we next expressed a step-function opsin with ultra-high light sensitivity (SOUL), which elicits prolonged but likely subthreshold depolarization after light stimulation, in E18-TRAPed neurons^69^. We then transcranially activated E18-TRAPed piriform neurons from P5–7 by placing an optic fiber over the scalp covering the piriform cortex (**Figure 6D**). Brief transcranial activation was sufficient to drive robust *Fos* expression in these neurons (**Figure S8B, C**). We then obtained whole-cell recordings from randomly selected layer 2 principal neurons and measured EPSCs evoked by electrical stimulation, as described above (**Figure 6D**).

Following chronic activation of E18-TRAPed neurons *in vivo*, we found significant increases in both recurrent (**Figure 6E**, left) and afferent (**Figure S8H**, left) EPSCs, suggesting that increasing activity of these E18-TRAPed piriform cells promotes recurrent and afferent connectivity. To again examine if these changes were specific to directly activating TRAPed neurons, we performed parallel experiments using *Ubc-CreER^T2^* to express SOUL in similar number of random piriform cells (**Figure S8D, E**). Chronic activation of this random subpopulation did not increase the amplitude of recurrent or afferent EPSCs (**Figure 6E, S8H**, right). These findings suggest that activating E18-TRAPed neuronal populations in early postnatal development promotes recurrent and afferent connectivity in the piriform network.

Next, we recorded spontaneous activity in pairs of randomly selected neurons. Chronic stimulation of TRAPed populations *in vivo* increased the frequency of spontaneous large and complex postsynaptic currents that were synchronized across simultaneously recorded neurons (**Figure 6F, G**). These events resemble giant depolarizing potentials (GDPs), which are thought to be population events in neonatal recurrent networks, such as neonatal hippocampus^31^. Indeed, pharmacological silencing of recurrent connections with baclofen completely abolished these events (**Figure 6G**). This further suggests that chronic activation of TRAPed cells increases piriform recurrent network. Once again, the increase in these large, synchronized events was specific to the activation of E18-TRAPed neurons as this was not observed following chronic stimulation of a random population of piriform neurons using *Ubc-CreER^T2^* mice (**Figure 6H**).

Finally, we examined whether transient activation of E18-TRAPed neurons during a neonatal period has long-lasting effect on piriform cortex connectivity. We transcranially activated SOUL-expressing E18-TRAPed piriform neurons from P5–12, and then performed whole-cell patch recording of piriform neurons in slices from P30 mice (**Figure 6I**). Again, both recurrent and afferent postsynaptic responses were significantly increased (**Figure 6J, S8I**), suggesting that the increased recurrent and afferent connectivity due to the transient increase in E18-TRAPed cells activity in neonates persist into young adulthood. Lastly, synchronized postsynaptic currents across simultaneously recorded neurons in P30 slices rarely occurred even after activation of E18-TRAPed neurons (**Figure 6K**), suggesting that this may be a transient property occurring during the maturation of recurrent connectivity in piriform cortex.

Taken together with our *in vivo* findings that TRAPed neurons have increased functional connectivity and lead population activity, these results suggest that E18-TRAPed piriform neurons are a hub-like population whose activity is necessary and sufficient to promote strengthening of recurrent connectivity in the developing piriform network (**Figure 2G_3_**).

## DISCUSSION

Here, our use of embryonic TRAP with whole-brain imaging provided the first comprehensive survey of embryonic neuronal activity in a mammalian brain *in vivo*. Previous observations of network activity and studies focusing on isolated cell types in early development have provided insight into activity-dependent phenomena in specific circuits^29,31,33,50,70^. Using TRAP, we identified many brain regions with prenatal activity worth further investigation, and focused on the embryonic and neonatal piriform cortex, a highly recurrently connected network in the mature brain. By tracking these neurons across development and manipulating their activity during neonatal stages, we found that embryonically active piriform neurons exhibit hub-like properties transiently during a developmental stage coinciding with recurrent circuit maturation (**Figure S2J**). The longitudinal tracking of this neuronal population afforded by TRAP provides direct evidence of the importance of such neurons specifically during early postnatal development. While previous studies have described the presence of hub neurons and how they affect network dynamics^32,33^, our manipulation of this functionally-defined embryonically active population demonstrated how their activity modulates the development of recurrent cortical connectivity.

The necessity of neonatal olfaction for survival and the importance of recurrent connectivity in adult piriform made the developing piriform cortex an attractive system for studying activity-dependent formation of recurrent circuits^18,19,40^. By establishing *in vivo* electrophysiological recordings of neonatal neuronal populations, we were able to study E18-TRAPed neurons within the general piriform network across different developmental stages. The high temporal resolution provided by our electrophysiological approach enabled identification of functional connectivity at monosynaptic time scales, allowing us to capture the hub-like functional connectivity of E18-TRAPed piriform neurons *in vivo*. Further, because our recordings were stable without anesthesia, we were able to investigate both sensory-driven and odor-independent network dynamics. These approaches enabled us to discover that E18-TRAPed cells lead spontaneous synchronized population activity (**Figure 5H**) without having a significant role in leading (**Figure S4E**) or amplifying odor responses (**Figure S5**) in neonatal piriform cortex.

A previous *ex vivo* study in the adult rat olfactory cortical slices described a “hierarchical architecture” in which a small population of densely interconnected neurons projects broadly to the rest of the network^71^. While the functional implication of this architecture *in vivo* has not been explored, our finding of an interconnected hub-like E18-TRAPed piriform population suggests that such an architecture may facilitate recurrent circuit formation. Reverberant excitation within this interconnected hub-like population and broad projection to the rest of piriform cortex may afford TRAPed neurons the ability to modulate synchronized population activity or general excitatory drive in the network. Hebbian plasticity in recurrent synapses during synchronized population activity or secondary to broadly increased excitation are potential mechanisms through which manipulation of TRAPed cell activity modulates recurrent connectivity strength (**Figure 6**). In addition to shaping the development of piriform architecture, this principle may be extended to other highly recurrently connected hippocampal circuits or intracortical connectivity in the neocortex.

Within the piriform cortex, our single-neuron-resolution analyses implicated the deep-layer piriform neurons as predominantly active in embryonic and neonatal stages (**Figure 2E, F**; **Figure S1F**) and as peak leaders of population activity (**Figure S6E**). This also provides insight into the a function of these deep neurons whose role in olfaction has been understudied compared to the more abundant layer 2 neurons^18,23^. Specifically, the fact that E18-TRAPed piriform neurons do not preferentially respond to neonatally-salient odors, respond similarly to maternal versus novel odors, and do not amplify odor responses, and instead have distinct features in odor-free epochs suggests a sensory-independent developmental phenomenon in this network. Although the piriform cortex has long-range recurrent connectivity compared to more local connectivity within neocortex, such hub-like neurons modulating sensory-independent network development in local microcircuits or long-range connections in other cortical and hippocampal regions may exist using similar developmental principles^45,72^.

The permanent labeling capacity of TRAP allowed us to study the embryonically active piriform population both in neonates and in adult mice. We were thus able to identify that the transience of this population’s hub-like properties coincided with a period of recurrent connectivity strengthening (**Figure S2J**). The general increase in piriform recurrent connectivity and low degree of inhibitory odor responses (**Figure 4F, S4K**) during this developmental stage may create ideal conditions for forming broad recurrent connectivity. Such a period may be followed by later stages of synaptic refinement to specify connectivity and establish the discrete ensemble representation of odors observed in mature piriform cortex^22,23,63,73^. In addition, recurrent connectivity strength at early stages may modulate the general degree of excitation in a network, which is known to strengthen developing sensory inputs by promoting dendritic NMDA-spikes^74^. This is consistent with our finding that activation of E18-TRAPed cells in early development also led to an increase in afferent input strength (**Figure S8H, I**). Further studies modulating hub-like populations at various ages and examining odor response dynamics in adult mice would provide insight into the different developmental stages of the piriform network. For example, persistent strengthening of recurrent connectivity, as was observed following E18-TRAPed cell stimulation, may promote odor representation stability across brain states in the adult^19^.

Despite the advantages of TRAP, there are several limitations of using a genetic approach to target active neurons. First, the direct relationship of activity and *Fos* expression is difficult to establish *in utero*. Second, TRAP is inherently limited to accessing neurons in developmental stages after the time of initial labeling, and thus does not afford study of embryonically active piriform neurons at the very time when their activity is TRAPed. However, the co-expression of other immediate early genes in embryonic *Fos*-expressing neurons, persistence of high activity in E18-TRAPed neurons postnatally, and the finding that E18-TRAPed neurons have hub-like properties in neonatal piriform, all suggest that this population has a distinct role in piriform development. While our study only investigated early postnatal stages, we hypothesize that similar hub-like properties of these neurons likely begin in late embryogenesis when they are TRAPed.

Lastly, because TRAP stochastically captures subsets of active neurons regardless of cell types, our approach resulted in an incomplete and mixed population that contain several distinct subpopulations of neurons (**Figure S7**). Thus, the activity of some subpopulations of neurons could regulate cell-autonomous development independent of circuit maturation (**Figure 2G_1_**), drive interconnectivity of co-active ensembles that either tune to neonatally-relevant odors or co-tune to other odors later in development (**Figure 2G_2_**), or be involved in other developmental processes. Nevertheless, despite the inability to capture all active neurons, our findings that the activity of E18-TRAPed neurons bidirectionally regulates recurrent interconnectivity suggests that at least a substantial subset of E18-TRAPed population does indeed act as a hub-like population in early development (**Figure 2G_3_**). Further studies with more granularly defined populations, perhaps using intersectional genetic approaches, may elucidate distinct roles of active subpopulations of embryonically active piriform neurons. Such work will likely reveal new mechanisms through which the dynamics of the developing brain shape the assembly of neural circuits.

## ACKNOWLEDGEMENTS

We thank Drs. Jun Ding, Carla Shatz, Regina M. Sullivan, Renzhi Yang, Fuu-jiun Hwang Sebastian H. Bitzenhofer, and Hiroaki Matsunami, as well as Robin Blazing for intellectual and technical guidance. We thank Dr. Guoping Feng for the SOUL Cre reporter mice and Dr. Martyn Goulding and Tomoko Velasquez for the TeNT Cre reporter mice. We thank Lucas Rivera-Encarnacion, Hannah Field, Daniel Pederick, Lijun Qi, Tom Hindmarsh Sten, Yunming Wu, and Alina Xiao and for critical comments on the manuscript. This work was supported by National Institutes of Health R01-NS050835 (L.L.) and R01-DC015525 (K.M.F.). D.C.W. is supported by a Stanford Bio-X fellowship. L.L. is an HHMI investigator.

## AUTHOR CONTRIBUTIONS

D.C.W., F.S.V., K.M.F., and L.L. designed the experiments. D.C.W. performed the experiments. D.C.W., F.S.V., and J.H.S. analyzed the data. D.C.W. and L.L wrote the manuscript with help from K.M.F. All the authors read and edited the final version of the manuscript.

## STAR*METHODS

### KEY RESOURCE TABLE

| <u>Reagent</u> | <u>Source</u> | <u>Identifier</u> |
| --- | --- | --- |
| **Antibodies and probes** |  |  |
| Rabbit Fos antibody | Synaptic Systems | Cat No. 226 008 |
| Rabbit RFP antibody | Rockland | Cat No. 600-401-379 |
| *Fos* probe | ACD Bio | Cat No. 316921 |
| *Arc* probe | ACD Bio | Cat No. 316911 |
| *Npas4* probe | ACD Bio | Cat No. 423431 |
| *tdTomato* probe | ACD Bio | Cat No. 317041 |
| *Vgat* probe | ACD Bio | Cat No. 319191 |
| **Experimental models: organisms/strains** |  |  |
| TRAP2 (Fos-CreER<sup>T2</sup>) | Jackson Laboratory | Stock Fos<sup>tm2.1(icre/ERT2)Luo/J</sup><br>Strain: 030323 |
| Ai14 (tdTomato Cre reporter) | Jackson Laboratory | Stock B6;129S6-Gt(ROSA)26Sor<sup>tm14(CAG-tdTomato)Hze/J</sup><br>Strain: 007908 |
| Ai32 (ChR2 Cre reporter) | Jackson Laboratory | Stock B6.Cg-Gt(ROSA)26Sor<sup>tm32(CAG-COP4*H134R/EYFP)Hze/J</sup><br>Strain: 024109 |
| TeNT Cre reporter | Dr. Martyn Goulding, Salk Institute | Stock B6;Gt(ROSA)26Sor<sup>tm2(GFP/tetX)Gld</sup><br>MGI ID: 3839913 |
| SOUL Cre reporter | Jackson Laboratory | Stock B6;129-Gt(ROSA)26Sor<sup>tm2(CAG-COP4*C128S*D156A*T159C,-tdTomato)Gfng/J</sup><br>Strain: 032301 |
| Ubc-CreER<sup>T2</sup> | Jackson Laboratory | Stock B6.Cg-Ndor1<sup>Tg(UBC-cre/ERT2)1Ejb/1J</sup><br>Strain: 007001 |
| CD1 WT | Charles River<br>Laboratory | Strain: 022 |
| **Chemicals, peptides, and recombinant proteins** |  |  |
| 4-hydroxytamoxifen | Sigma | Cat No. H6278 |
| Picrotoxin | Tocris | Cat No. 1128 |
| Baclofen | Tocris | Cat No. 0417 |
| CGP 55845 | Tocris | Cat No. 1248 |
| Ethyl butyrate | Aldrich | Cat No. E15701 |
| Hexanal | Aldrich | Cat No. 115606 |
| Ethyl acetate | Sigma | Cat No. 34858 |
| Isoamyl acetate | Tokyo<br>Chemical<br>Industry | Cat No. A0033 |
| Garlic oil | Sigma | Cat No. W250320 |
| Vanillin | Sigma | Cat No. V1104 |
| **Critical commercial assays** |  |  |
| RNAscope Multiplex<br>Fluorescent Reagent<br>Kit V2 | ACD Bio | http://acdbio.com |
| **Software and algorithms** |  |  |
| ClearMAP | Marc Tessier-Lavigne<br>Laboratory | http://idisco.info/clearmap-2/ |
| ImageJ | NIH | http://imagej.nih.gov/ij/ |
| GraphPad Prism 9 | GraphPad<br>Software | http://www.graphpad.com |
| pClamp 11 | Molecular<br>Devices | http://www.moleculardevices.com/ |
| MATLAB | Mathworks | http://www.mathworks.com |
| Python 3.8.5 | Python<br>Software<br>Foundation | http://www.python.com |

#### Lead contact

Further information and requests for resources and reagents should be directed to and will be fulfilled by the lead contact, Liqun Luo.

#### Materials availability

This study did not generate new unique reagents.

#### Data and code availability

Data and custom MATLAB code used to generate all analyses are available from the Lead Contact upon request.

### EXPERIMENTAL MODELS AND SUBJECT DETAILS

#### Animals

All animal procedures followed animal care guidelines approved by Stanford University’s Administrative Panel on Laboratory Animal Care and Duke University’s Institutional Animal Care and Use Committee.

To generate *TRAP2;Ai14* and *TRAP2;Ai32* mice, we crossed *Fos2A-iCreER* mice (FosTRAP2, Jackson Laboratory, Stock 030323)^36^ with the tdTomato Cre reporter mice (*Ai14*, Jackson Laboratory, Stock 007914)^47^ or ChR2-eYFP Cre reporter mice (*Ai32*, Jackson Laboratory, Stock 012569)^47^. These mice were subsequently backcrossed to wild-type CD1 mice for two generations, then bred to homozygosity in all involved alleles, resulting in *TRAP2/TRAP2;Ai14/Ai14* or *TRAP2/TRAP2;Ai32/Ai32* mice. Backcrossing to CD1 improved tolerance and litter size after tamoxifen administration to pregnant dams. Slice electrophysiology experiments were performed from a cross of *TRAP2;Ai14* and *TRAP2;Ai32* mice, resulting in *TRAP2/TRAP2;Ai14/Ai32* mice.

*Ubc-CreER^T2^/+* (Jackson Laboratory, Stock 007001)^65^ mice were backcrossed to wild-type CD1 for two generations before use in experiments. To generate *TRAP2;SOUL*, *Ubc-CreER^T2^;SOUL* mice, *TRAP2;LSL-TeNT*, or *Ubc-CreER^T2^;LSL-TeNT* mice, *SOUL-P2A-tdT* Cre reporter mice (*SOUL*, Jackson Laboratory 032301) or *LSL-TeNT* (*TeNT*, obtained from Dr. Martyn Goulding, Salk Institute)^76^ males were crossed to *TRAP2/TRAP2* or *Ubc-CreER^T2^/+* females.

### METHOD DETAILS

#### Tamoxifen preparation, administration, and neonatal mouse fostering post-administration

4-hydroxytamoxifen (4OHT; Sigma H6278) was dissolved at 20 mg/ml in 100% ethanol by shaking at 37°C for 1hr and was then aliquoted and stored at –20°C for up to 3 months. Before use, 4OHT was redissolved in ethanol by shaking at 37°C for 30 min, a 1:4 mixture of castor oil:sunflower seed oil (Sigma 259853 and Sigma S5007) was added to give a final concentration of 10 mg/ml 4OHT, and the ethanol was evaporated by vacuum centrifugation. The final 10 mg/ml 4OHT solutions were used within several hours after they were prepared.

For postnatal 4OHT administration, intraperitoneal (IP) injections were performed at a dosage of 50mg/kg of animal body weight. For 4OHT applications targeting embryonic brains, oral gavages (OGs) were performed on the pregnant dam at E18 (where E0.5 is defined as the morning when the mucus plug was found, indicating copulation had occurred the night prior). OGs were done at concentrations of 50mg/kg of animal body weight (for histology and iDISCO in *TRAP2/+;Ai14/+* mice) or 150mg/kg of animal body weight (for slice electrophysiology in *TRAP2/TRAP2;Ai14/Ai32* mice, and *in vivo* electrophysiology in *TRAP2/TRAP2;Ai32/Ai32* mice). The increase in 4OHT dose was to compensate for the relatively low recombination efficiency with TRAP and the *Ai32* reporter allele (**Figure S2A, B**). For SOUL and TeNT experiments, 200mg/kg 4OHT was used to increase labeling density as these mice were heterozygous for both alleles. Pups born after E18 4OHT administration were given to a foster mother (either a WT CD1 dam or a dam of the same genotype, who had been fed the same diet and had given birth to a litter the same day or one day prior) as perinatal 4OHT administration typically reduces or eliminates milk production or letdown in the dam.

#### iDISCO+ and whole-brain imaging

To prepare brains for the iDISCO+, lightsheet imaging, and automated cell counting, adult (>P40) *TRAP2;Ai14* animals were perfused transcardially with PBS and 4% PFA, and post-fixed overnight at 4°C in 4% PFA. Samples were then processed with the iDISCO+ immunolabeling protocol, as detailed at http://www.idisco.info. Samples were stained with an RFP primary antibody (Rockland 600-401-379) at 1:500, followed by an Alexafluor-647 secondary antibody (ThermoFisher Scientific) at 1:500.

After clearing, samples were submerged in dibenzyl ether and imaged on a light-sheet microscope (Ultramicroscope II, LaVision Biotec). Samples were scanned with a step-size of 3 µm using the continuous light-sheet scanning method with the included contrast adaptive algorithm for the 640 nm channel (20 acquisitions per plane), and without horizontal scanning for the 488 nm autofluorescence channel.

#### Histology

To collect brains for histology, animals were perfused transcardially with PBS and 4% PFA, and post-fixed overnight at 4°C in 4% PFA. Free floating coronal sections were cut at 60-µm thickness using a Leica Vibratome system.

Immunostaining was performed by incubating sections in primary antibodies for c-Fos (Synaptic Systems 226 008) or tdTomato (Rockland 600-401-379) in PBST (phosphate-buffered saline + 0.1% Triton) overnight at room temperature, washed 3x with PBST, secondary antibodies in PBST overnight at room temperature, washed 3x with PBST, and mounted in Vectashield (Vectorlabs).

#### RNAscope

RNAscope was performed according to the manufacturer’s instruction manual using standard probes designed by ACDBio (*Fos,* Cat No 316921; *Arc,* Cat No 316911; *tdTomato,* Cat No 317041; *Vgat,* Cat No 319191). Changes made to accommodate embryonic and neonatal tissue include the following:

Fresh frozen coronal sections were cut at 20-µm thickness using a Leica Cryostat system. Slides were dried overnight at –20°C, and sections were post-fixed with 4% PFA for 2 hr at room temperature. Sections were treated with H_2_O_2_ for 10 min at room temperature, followed by Protease III for 10 min at room temperature.

#### Electrophysiology in brain slices

Neonatal brain slices were obtained from *TRAP2/TRAP2;Ai14/Ai32* (high-density TRAP regime) *TRAP2/+;Ai14/Ai32* (low-density TRAP regime), *TRAP2/+;SOUL/+*, or *TRAP2/+;TeNT/+* animals. Brains were removed and placed in ice-cold carbogenated slicing artificial cerebrospinal fluid (ACSF) that contained (in mM) 83 NaCl, 2.5 KCl, 1 NaH_2_PO_4_, 26.2 NaHCO_3_, 22 glucose, 72 sucrose, 0.5 CaCl_2_, and 3.3 MgSO_4_, at 305 mOsm. Coronal sections (300 μm) were sliced on a Leica vibratome in this same ACSF. Slices were allowed to recover at 32°C for 45 min and then at room temperature for 20 min before recording. Slices were then placed in room temperature carbogenated recording ACSF that contained (in mM) 119 NaCl, 2.5 KCl, 26 NaHCO_3_, 1 NaH_2_PO_4_, 1.5 MgSO4, 2.5 CaCl_2_, and 11 glucose. Signals were recorded with a 5× gain, low-pass filtered at 2 kHz, digitized at 10 kHz (Molecular Devices Multiclamp 700B), and analyzed with pClamp 10 (Molecular Devices). Whole-cell recordings were made using 4-MΩ pipettes filled with an internal solution that contained (in mM) 130 cesium gluconate, 5 NaCl, 10 EGTA, 10 HEPES, 4 MgATP, 0.3 NaGTP, 12 phosphocreatine, 5 QX-314, 10 TEA, at pH 7.3 and 295 mOsm. Alexafluor 555 hydrazide (Thermo Fisher A20501MP) was added to the internal solution to facilitate morphological analysis of recorded cells. Series resistance (R_s_) and input resistance (R_in_) were monitored throughout the experiment by measuring the capacitive transient and steady-state deflection, respectively, in response to a −5 mV test pulse.

Cells were clamped at −70 mV to measure EPSCs. Miniature EPSCs (mEPSCs) were recorded over 5 min after breaking into the cell in whole-cell configuration and achieving stable voltage clamp in ACSF containing 1 µM TTX (Tocris 1069).

Optogenetic stimulation responses were evoked by a 200-µm diameter 0.39 NA flat cleave patch cable (Thor Labs M136L02) placed 200 µm lateral to and 500 µm above the recorded cells. Laser power was 10 mW (this high laser power was required to obtain responses in most neonatal brain slices, potentially because ChR2 expression is low and synapses are immature).

Electrical stimulation responses were evoked by a concentric bipolar microelectrode (FHC) placed in piriform cortex layer 3 approximately 250 µm away from the recorded cells. To find the stimulus amplitude required to stimulate all local axon terminals, we created stimulus amplitude-response curves to find the stimulation amplitude required to saturate the evoked synaptic current amplitude. For neonatal piriform neurons of excitatory morphology, this was typically 100–150 µA. Recordings of neurons thus consisted of 5 sweeps, each typically containing one 200-µA 200-µs (although could require up to 400-µA 400-µs) stimulus followed by a 45-s rest interval to allow the cell to recover.

To isolate recurrent input in piriform cortex, we perfused ACSF containing 50-µM baclofen (Tocris 0417) and allowed the cell to stabilize for 10 min. To confirm that baclofen did not result in rundown of the cell, we subsequently perfused ACSF containing 10 µM CGP 55845 hydrochloride (Tocris 1248) to a subset of recorded cells to ensure that the pre-baclofen synaptic currents could be recovered.

#### SOUL chronic activation in neonatal mice and electrophysiology

Following E18-TRAP in *TRAP2/+;SOUL/+* or *Ubc-CreER^T2^/+;SOUL/+* mice, the TRAPed litter was combined with a nonTRAPed litter of the same genotype and given to the foster mother who did not receive a 4OHT administration. These litters were culled to a maximum of 8 pups. Beginning at P5 or P6, each pup was transcranially optically stimulated for 30 s using 10-mW 488-nm laser by placing a bare optic fiber over the skin covering piriform cortex. No surgery was performed on these animals. Only one hemisphere was stimulated, and the fiber was moved along the anterior-posterior axis to stimulate cells spanning the entire piriform cortex. Each pup was stimulated 4x / day from starting at P5. Both TRAPed pups and littermates who did not receive 4OHT received the same transcranial stimulation, and pup identity was blind to the experimenter. Following stimulation, pups were immediately returned to homecage.

Electrophysiology was performed at P7-9 for neonatal recordings or ∼P30 for young adult recordings. Acute slices were obtained as described in above sections, but in a room only illuminated with orange light to prevent activation of SOUL. To ensure SOUL is silenced during recording, slices were placed on the recording rig and illuminated with 10-mW 589-nm light from a bare optic fiber placed approximately 1 mm above the slice within the ACSF bath immediately before recording evoked postsynaptic currents. Dual-patch whole-cell recordings was established in two neighboring superficial pyramidal cells. Cells were in ACSF containing 100 µm picrotoxin (PTX, Tocris 1128) and clamped at −70 mV to measure excitatory synaptic currents. Electrical stimulation and baclofen application were performed as described in above sections.

#### Neonatal mouse surgery for Neuropixels recordings

To prepare neonatal mice for Neuropixels recordings, P6–10 mice were deeply anesthetized with isoflurane. Neonatal mice were stabilized on the stereotax using custom 3D-printed head holders, which could be attached onto standard earbars. The scalp was removed, lidocaine was applied locally to the cut tissue, and craniotomies were made unilaterally above APC and PPC. Coordinates were measured from bregma, and APC and PPC coordinates were scaled down from adult coordinates (APC: AP 1.2, ML 2.1; PPC: AP –0.82, ML 3.75; all in millimeters). Scaled coordinates were calculated as the following: AP scaling factor = (pup bregma-to-lambda distance)/4.2, ML scaling factor = (pup midline-to-skull-edge distance)/4.6. A 100-µm diameter craniotomy was made directly above these sites for Neuropixels access, and a separate 500-µm diameter craniotomy was made 400–500 µm medial or lateral to this site for optic fiber access. KwikCast (WPI) was used to cover the craniotomies. A headpost made from the needle tip of a flattened 18G needle placed over the hindbrain, and a groundpin composed of a silver wire and gold pin were placed into the hindbrain. Metabond (Parkell) was applied broadly to cement the headpost and groundpin. Neonatal mice were then removed from anesthesia allowed to recover under a heat lamp (body temperature maintained at 32°C) for 3–4hr, then briefly held 1 cm from its native dam and allowed to feed for 1–2 min. Only mice able to locate the dam when held 1 cm away were used in recordings. Mice were subsequently fed 50 µl of warmed half & half (Organic Valley) before being placed on the recording rig.

Prior to P6, the skulls of neonatal mice were not stable enough to keep a headfixed mouse from moving, and thus significant drift would occur throughout the recording, especially due to mouse movement in response to odor. Because of this, only mice P6 or older were used for recording.

#### Neuropixels recording

Neonatal mice were awake and unanesthetized for the duration of the recording. Mice were headfixed and placed on a custom 3-D printed chamber to allow for a more natural head angle during recording and kept at 32°C with a heatpad. Each mouse was only recorded in one session (specifically, at 3–4hr after the headpost surgery and craniotomy). After recording, mice were perfused for histological confirmation of successful probe insertion into the piriform cortex. Each mouse had two electrodes implanted simultaneously (one in APC, one in PPC).

*In vivo* electrophysiological recordings of awake P6–10 mice were performed using Phase 3B Neuropixels electrodes with 384 active recording sites along the tip of the probe. Kwikcast above the craniotomy was removed and the craniotomy was filled with 0.9% NaCl in water. Electrodes were coated with a red lipophilic dye (DiI, Thermo Fisher) and dried for 3–5 min. Electrodes were then lowered slowly (2–3 μm/s) into brain using a micromanipulator (Scientifica PatchStar) until the electrode bent, indicating that it had reached the bottom of the skull, after which the electrode were retracted 100 µm. Typically, the tip of the electrode had a depth (DV) of ∼4 mm at APC and ∼4.5 mm at PPC of a P6–10 pup. Electrodes were then allowed to settle in the brain for 45 min before recording. Recordings were sampled at 30 kHz amplified (500x gain) and bandpass filtered (0.3–10 kHz). Signals were digitized with a CMOS amplifier and multiplexer on the Neuropixels electrode, then written to the disk using SpikeGLX software. Respiration was measured with a microbridge mass airflow sensor (Honeywell AWM3300V) positioned directly opposite the animal’s nose and signals were recorded using a MCC DAQ board. Custom code was written for the Open Ephys GUI to control the olfactometer and synchronize it with the MCC data stream online. Neuropixels and MCC data were synchronized offline.

To optotag TRAPed units in *TRAP2/TRAP2;Ai32/Ai32* mice, we attached a 400-µm 0.39-NA optic fiber (Thor Labs) to each Neuropixels probe, approximately 500 µm lateral to the probe with the fiber tip terminating ∼1.5 mm from the tip of the electrode. A 470-nm LED (∼15 mW at the fiber tip) was used to present brief pulses at the beginning and end of each recording (30 pulses of 20 ms at 0.5 Hz), the responses of which were later used to determine optotagged units. Units that had significant increases in action potentials within ∼10 ms in neonatal mice and ∼5 ms in adult mice in response to >95% of the pulses were considered optotagged units. Nearly no neonatal units were optotagged within 5 ms (this difference in latency may be due various immature electrophysiological properties of neonatal neurons, as well as lower expression of ChR2 in P6-10 mice, leading to decreased efficacy in light induction of spikes). Using a ∼10 ms window in neonates and ∼5 ms in adults led to roughly similar proportions of optotagged units (2-4% of recorded units).

Each recording session began by recording of spontaneous activity for 15 minutes without any optogenetic or odor stimuli. A 2-min optotagging period followed the spontaneous epoch before odor presentation began. Odorants were then presented in a specific order, with a 2 second odor stimulus with a ∼22 second interstimulus interval. This series was repeated so that each odorant was presented 10 times. After the last odorant presentation, an additional 2-minute optotagging period was carried out to ensure the stability of the optotagged units. Histology was performed after each recording; only units confirmed to be within piriform cortex were used for further analysis.

#### Odorants and odorant delivery

Monomolecular odorants were diluted to 0.02% in mineral oil (this low concentration was determined by calibrating the concentrations to evoke similar numbers of spikes across monomolecular and mouse odors in initial trials). The odorant set consisted of hexanal (Aldrich 115606), ethyl butyrate (Aldrich E15701), ethyl acetate (Sigma 34858), and isoamyl acetate (Tokyo Chemical Industry, A0033). Monomolecular odorants were presented by passing air through odorants contained in 20-ml amber vials containing 5 ml of odorant solution.

Mouse odorants were collected directly from dams with signature odors, or male adult mice. To create dams with signature odors, pregnant dams were fed water containing garlic oil (Sigma W250320, diluted to 0.05% v/v, and shaken at 37°C overnight) or vanillin (Sigma V1104, 75 mM) beginning when embryos were at the E14 stage. Ventra odors consisted of air flowed directly over an anesthetized mouse ventrum, using custom 2 cm x 2 cm 3D-printed domes designed to pump in, mix, and capture air over the ventra during the *in vivo* recording sessions. Mouse milk was collected from anesthetized dams (anesthetized with 100 mg/kg ketamine, 10 mg/kg xyzaline, and with 5 IU of oxytocin to promote milk letdown). Mouse urine was collected directly from scruffed mice. Amniotic fluid was collected at E17.5 using an 18G syringe to extract amniotic fluid from C-sectioned uteri. Mouse milk, urine, and amniotic fluid were snap frozen with liquid nitrogen and stored at –80°C until use. Liquid mouse odorants were presented by passing air through odorants contained in a 2-ml screw cap tube containing 200 µl of mouse odorant. Odorants were delivered using a custom-built olfactometer controlled by custom MATLAB scripts, the construction of which was previously described^21^.

#### Optogenetic activation of TRAPed cells during odor presentation

To activate TRAPed populations during odor delivery, the same optical fiber from a 470-nm LED used for optotagging was used. Laser power was titrated by presenting 3x 300ms 0.25 Hz light pulses at 15 mW (full power), 7.5 mW, 4.5 mW, 1.5 mW, and 0.15 mW at the beginning of each experiment. Near-threshold laser power was determined as the average of the lowest laser power that elicited spikes and the highest laser power that did not elicit spikes in most TRAPed units. Odor presentation with optical stimulation was performed at the end of each experiment, after the regular odor presentation series was complete. During a 1 second odor delivery, continuous optical stimulation was activated at near-threshold laser power. Odor delivery was done as described above, and subsequent analysis was done the same as for odorant responses as described below.

### QUANTIFICATION AND STATISTICAL ANALYSES

#### Automated cell counting of iDISCO samples and cell type quantification

To quantify iDISCO+ samples, we re-trained the TrailMap model^77^ using sample images from *TRAP2;Ai14* mouse brains to automatically segment tdTomato+ cells by neuronal morphology. After segmentation, the 3-D coordinates of labeled soma were extracted using customized codes written in MATLAB and were registered to the Allen Mouse Brain CCFv3 using Elastix (Klein et al., 2010).

To quantify cell types within piriform cortex, 150–300 µm optical slabs were taken from whole-brain imaging samples in order to more clearly visualize cellular morphology, and manually counted using previously commonly reported cell types as a reference^22,58,60^.

#### Electrophysiology in brain slices

Processing of raw electrophysiological data and spontaneous postsynaptic current quantification were performed with the ClampFit Advanced Analysis Module. For optogenetically evoked synaptic currents, responses were measured as the peak current amplitude of the first peak within the first 10 ms. For electrical stimulation of all inputs, evoked synaptic current responses were measured as the amplitude of the first peak within the first 10 ms. Analysis was based on the average of 3–5 sweeps.

After baclofen was added to ACSF to silence recurrent inputs, synaptic currents from recurrent inputs were calculated as the difference in optogenetic or electrical stimulation–evoked synaptic currents between the without- and with-baclofen conditions.

For electrophysiology experiments following SOUL stimulation, synchronized event frequency was calculated as the number of events (synchronized between two simultaneously recorded neurons, surpassing 50 pA in magnitude and lasting more than 200 ms before returning to baseline) during a recording of spontaneous activity 3 min after establishing whole-cell recording configuration.

#### Neuropixels spike sorting

Spike sorting was performed offline using Kilosort2.5 with default parameters. The clusters were manually curated using Phy2 to determine whether they were “Good” (well-isolated and stable) or “MUA” (poorly isolated or unstable units). All units classified by Kilosort as “Good” were manually inspected, and clusters with high signal-to-noise ratio, clean spike time auto-correlograms were kept for analysis.

#### Neuropixels data analysis

To quantify odorant response dynamics, we computed a smooth kernel density function (KDF) using a 20-ms Gaussian kernel (Chronux toolbox psth routine) for each cell-odor pair. Response indices were calculated as briefly described^21^. Briefly, the reliability of differences between a cell’s spike counts within a 2-s odor presentation window vs 2-s baseline prior to odor presentation was quantified by computing the area under the receiver operating characteristic curve (auROC). The response index was obtained by multiplying the auROC by 2 and subtracting 1. Peak latency was the time of the maximum of a KDFs were computed from spikes within the 2-s response window, and response duration was the width at 50% of the maximum of this peak.

Lifetime sparseness is an index of the breadth of odor tuning^78^, and is calculated from the distribution of spikes in response to different odors for a given unit, and is calculated as S_L_ = {1−[(Σr_i_/n)^2^/Σ(r_i_^2^/n)]}/[1−(1/n)], where r_i_ is the trial-averaged response to the i-th odor and n is the number of odors.

A custom MATLAB script was written to analyze spontaneous activity of the population and of each unit within a 15-min spontaneous epoch prior to odor presentation. Firing rates were calculated over 10-ms time bins, and a Z-score for each time bin was calculated relative to the average firing rate across the spontaneous epoch. Average population activity was calculated by taking the mean Z-score across all units. Peaks in population activity were calculated using MATLAB’s ‘findpeaks’ function on smoothed population Z-scores (using a moving average filter of 20 time bins). Population events were defined as peaks in which the population Z-score was one standard deviation above the median Z-score and over 30% of the recorded units also had an individual unit peak within the width at 50% of the maximum Z-score of the respective population peak. The 30% criterion was used to exclude peaks in population Z-score that were a result of a small number of units with transient extremely high firing rate, which was not the population level event we sought to quantify. These parameters were determined manually in a manner blind to all recording conditions (e.g., unit ID, age, depth of unit, etc.), and several other sets of parameters were tested with similar results. Individual unit peaks were calculated using MATLAB’s ‘findpeaks’ on smooth unit Z-scores (using a moving average filter of 20 time bins), and only peaks with Z-score one standard deviation above the median Z-score were kept for analysis. Individual units were considered to participate in population peaks if the individual unit peak fell within the width at 50% of the maximum Z-score of the population peak. Latency to peak was calculated as the average time difference between an individual unit’s peaks and the respective population peaks around which that unit peak occurred.

Functional connectivity *in vivo* was determined using a custom C++ script creating correlograms of all pairs of units within a single recording. Significance of functional connectivity within synaptic latencies was determined using previously described jitter methods^79^. Functional connections for each unit included significant correlations from spontaneous and odor-presentation epochs.

A depth index was determined for each recorded unit from a combination of Neuropixels and histology data. The position of each layer relative to the tip of the probe was determined by post-recording histology. The position of each recorded unit along the probe tract was determined by the distance of the channel (corresponding to the largest amplitude of the unit’s waveform) from the tip of the probe. A depth index was then calculated to account for the difference in layer positioning and thickness in each probe insertion, and for the fact that the dense layer 2 was significantly thinner than the large and sparse layer 3. Depth indices were calculated as follows: units from the cortical surface to the beginning of layer 2 were given a score of 0 to 2, depending on their position along the cortical surface−layer 2 axis. Similarly, units within layer 2 were given a score of 2 to 4, and units from the beginning of layer 3 to the end of piriform were given a score of 4 to 8. Data are presented with “cortical surface” corresponding to a depth index of 0, “L2-L3 boundary” to 4, and “end of piriform” to 8.

Bootstrapped distributions were created when comparing distributions of TRAPed versus all cells (for lifetime sparseness, firing rate, odor response dynamics, functional connectivity, participation in population peaks) to account for the different number of units in each population (n_TRAP_ = 63, n_All_ = 2020 units in neonatal piriform; n_TRAP_ = 17, n_All_ = 864 units in adult piriform). An estimated sampling distribution for all cells was created by taking the mean of the metric in question of a randomly sampled 63 units from the total population of 2020 units for neonate or 17 units from the total population of 864 units for adult without replacement 1000 times for comparing TRAPed units to all units, or 1000000 times for comparing units within a cluster of interest to units in all other clusters (**Figure S7**). The p-value was obtained by comparing the mean of the metric in question for the TRAPed population to this estimated sampling distribution with α = 0.05, or with an adjusted α when performing Bonferroni correction for multiple comparisons.

#### Clustering of cell types *in vivo* by anatomical and physiological properties

To cluster piriform cell types from *in vivo* properties, 564 piriform units from 5 neonatal mice and 864 piriform units from 4 adult mice were included. These 9 mice had all features recorded, and neonatal mice that did not have certain odorants presented or did not have a spontaneous activity recording epoch were excluded.

The 30 features used were the following: age, anterior or posterior piriform cortex, individual response indices to 14 odorants (including mineral oil), average response index, peak odor response firing rate, latency to odor response peak, width of odor response peak, depth index, average delay to population peak, unit peak during population peak, lifetime sparseness, average firing rate, leading excitatory functional connectivity, lagging excitatory functional connectivity, leading inhibitory functional connectivity, lagging inhibitory functional connectivity, and total excitatory functional connectivity.

Clustering analysis was performed by a custom MATLAB script. Values for each feature were entered into columns of a matrix, and the columns were then normalized. PCA was used to reduce dimensionality to five PCs, determined by inflection point on an elbow plot. K-means clustering was then performed to assign each unit to one of five clusters (optimal number of clusters also determined by inflection point on an elbow plot). UMAP was used to generate a 2D projection of clusters. Each cluster was then compared to the collection of all other clusters along all 30 features, totaling 150 comparisons. TRAPed cell proportion within each cluster was determined by dividing the number of TRAPed cells within a cluster by the total number of cells within the corresponding cluster.

---

**Figure S1.**
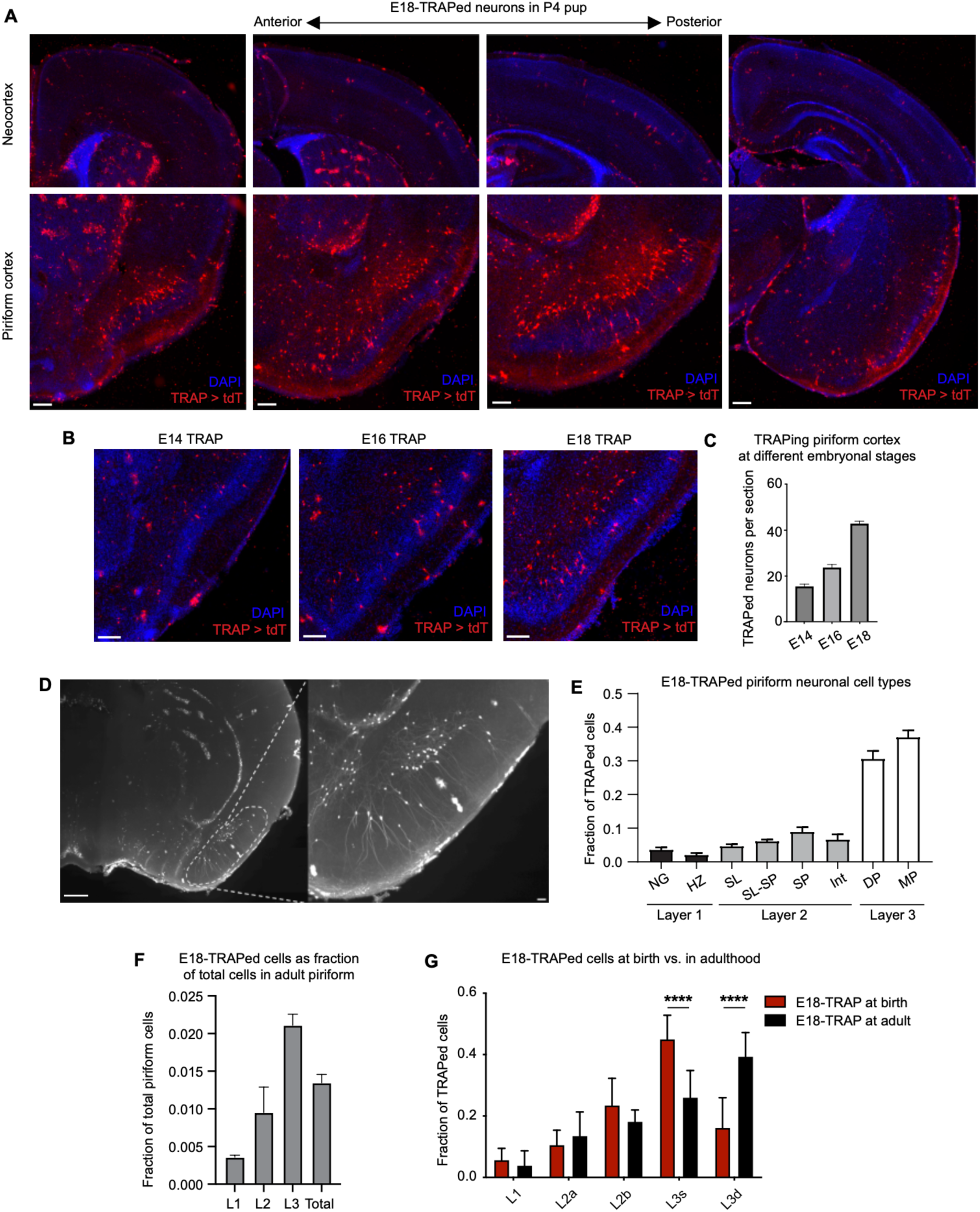
Additional characterization of embryonically TRAPed neurons, related to Figures 1 & 2. (A) Example images of E18-TRAPed neurons in neocortex and piriform cortex along the anterior-posterior axis from P4 pups. Note these animals were TRAPed with 150 mg/kg 4OHT (3x the dose used in experiments in **Figures 1, 2, and S1D–G**) to increase sensitivity of labeling. Scale bar, 100 μm (B, C) Example 60-µm sections (B) and quantification (C) of embryonically TRAPed neurons in piriform cortex when 4OHT were injected to label active neurons at E14, E16, and E18, respectively (n = 14–16 sections across 3 animals per age). Scale bar, 100 μm. (D) Left, a 300-µm coronal slab of a portion of a cleared P40 mouse brain TRAPed at E18. Right, higher magnification demonstrating a variety of TRAPed piriform neuron cell types. Scale bar, 100 μm. (E) Quantification of TRAPed neuron cell types within piriform cortex (n = 3 mice). Error bars, SEM. NG, neurogliaform; HZ, horizontal; SL, semilunar; SP, superficial pyramidal; SL-SP, intermediate between SL and SP; Int, interneuron; DP, deep pyramidal; MP, multipolar. (F) Layer distribution of E18-TRAPed cells in piriform cortex as a fraction of total cells in piriform cortex (n = 6 sections across 3 animals). Error bars, SEM. (G) Layer distribution of E18-TRAPed cells at birth (RNAscope for TRAP-induced tdTomato on brains from C-sectioned embryos at E19) or in adulthood (immunostaining on >P40 brains). Layers 2a (superficial layer 2), 2b (deep layer 2), 3s (superficial layer 3), 3d (deep layer 3) (n_E18, adult_ = 4 mice, n_E18, birth_ = 6 mice; ****p<0.0001, two-way ANOVA).

**Figure S2.**
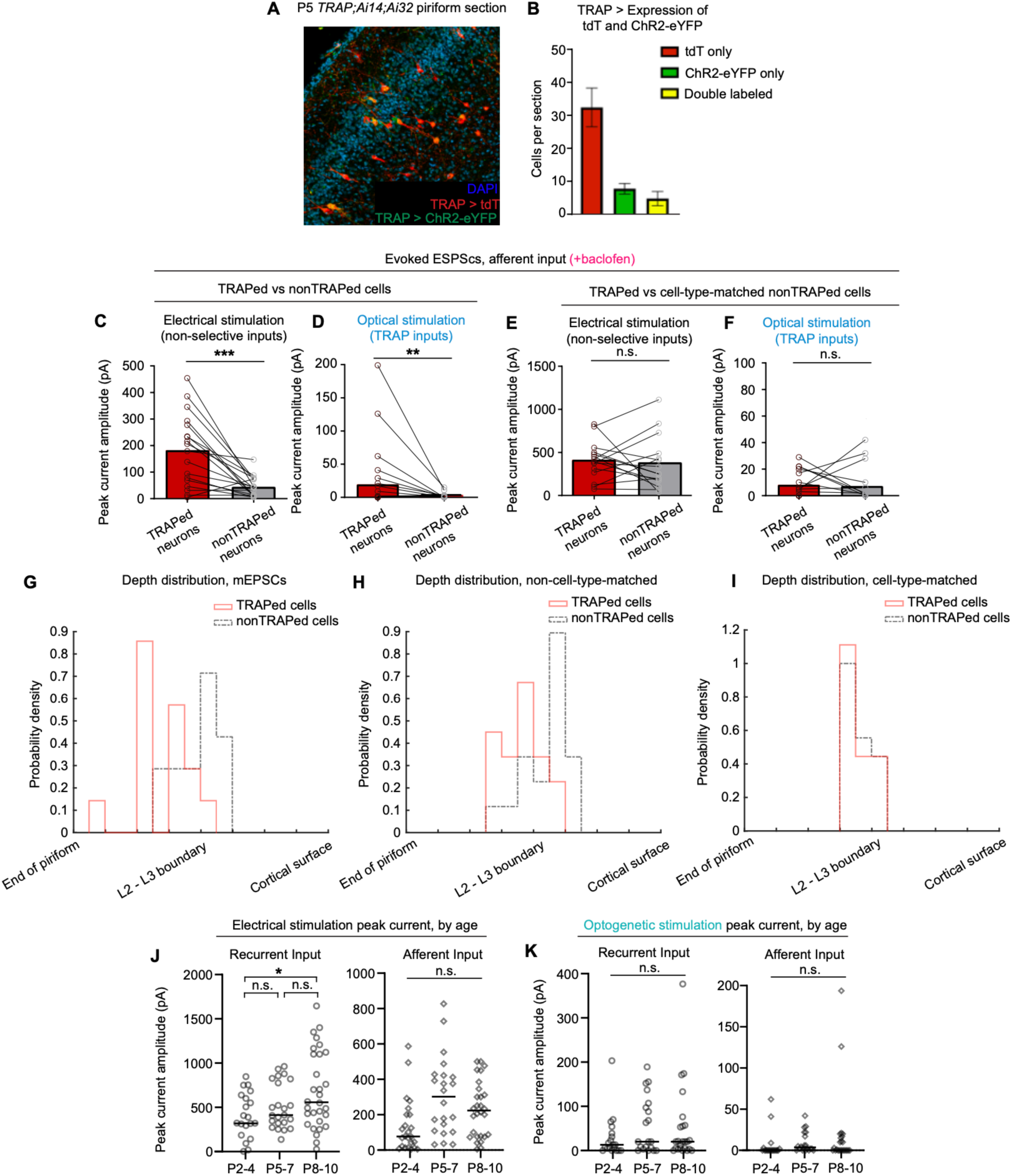
Additional physiological features of TRAPed neurons and recurrent piriform connectivity during early development, related to Figure 2. (A, B) Example (A) and quantification (B) of tdTomato- and ChR2-labeling in embryonic-TRAPed cells (n = 6 sections) (C, D) Summary of peak current amplitude under electrical stimulation of all afferent inputs (C) and optogenetic stimulation of TRAPed afferent inputs (D), using baclofen to silence recurrent axonal terminals. n = 18 per condition across 5 mice; *p<0.05, **p<0.005, ***p<0.001, paired two-tailed t-test (E, F) Summary of peak current amplitude under electrical stimulation of all afferent inputs (E) and optogenetic stimulation of TRAPed afferent inputs (F) to TRAPed vs cell-type-matched nonTRAPed neurons, using baclofen to silence recurrent axonal terminals. n = 17 per condition across 12 mice; *p<0.05, **p<0.005, ***p<0.001, paired two-tailed t-test (G–I) Layer distribution of cells in mEPSC and evoked EPSC experiments. (J, K) Evoked postsynaptic current of recurrent or afferent inputs from electrical stimulation of all axons (J) or optogenetic stimulation of TRAPed neuron axons (K), stratified by age of the animal of the recorded cell. n_P2-4_ = 29, n_P5-7_ = 23, n_P8-10_ = 30 for electrical, n_P2-4_ = 19, n_P5-7_ = 21, n_P8-10_ = 25 for optogenetic, across 19 neonatal mice; *p<0.05, one-way ANOVA with post-hoc Tukey HSD.

**Figure S3.**
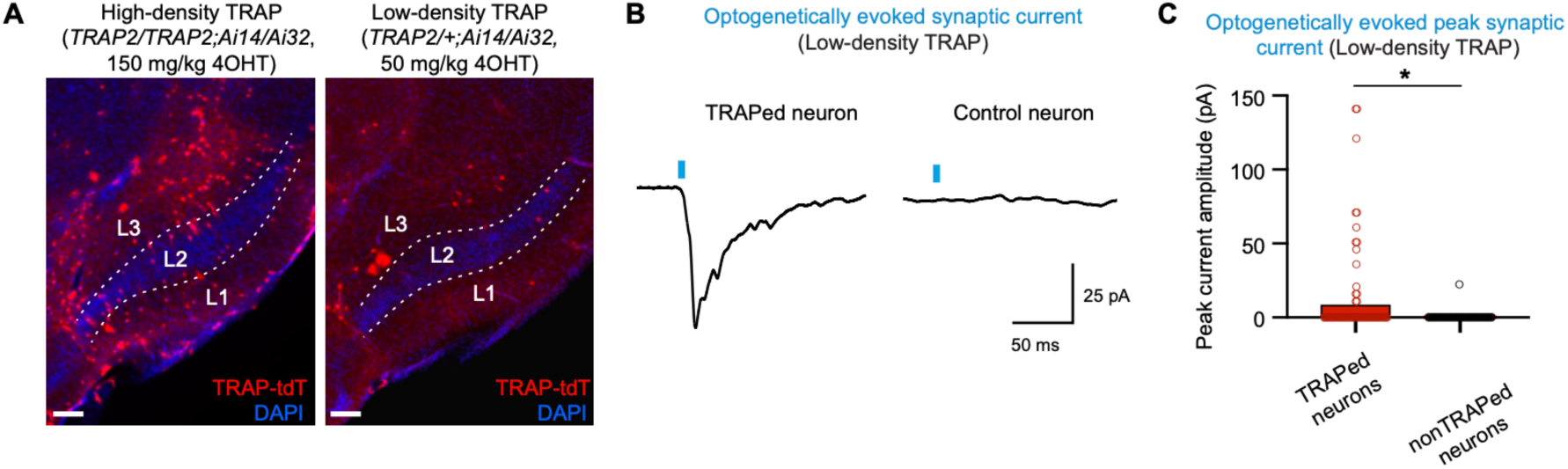
Synaptic connectivity of E18-TRAPed neurons in sparsely labeled *TRAP2;Ai14/Ai32* mice, related to Figure 2. (A) Example images of E18-TRAPed neurons in piriform cortex under high-density (high 4OHT dose, *TRAP2* homozygous) or low-density (low 4OHT dose, *TRAP2* heterozygous) TRAPing regimes. Scale bar, 100 μm. (B) Under the low-density TRAPing regime, example averaged trace of 5 evoked postsynaptic currents in a TRAPed vs a control cell (both clamped at –70 mV), following optogenetic stimulation of synaptic inputs from TRAPed neurons. (C) Under the low-density TRAPing regime, summary of evoked postsynaptic current under optogenetic stimulation of inputs. n = 68 for TRAPed neurons, n = 42 for control neurons, *p<0.05, unpaired t-test.

**Figure S4.**
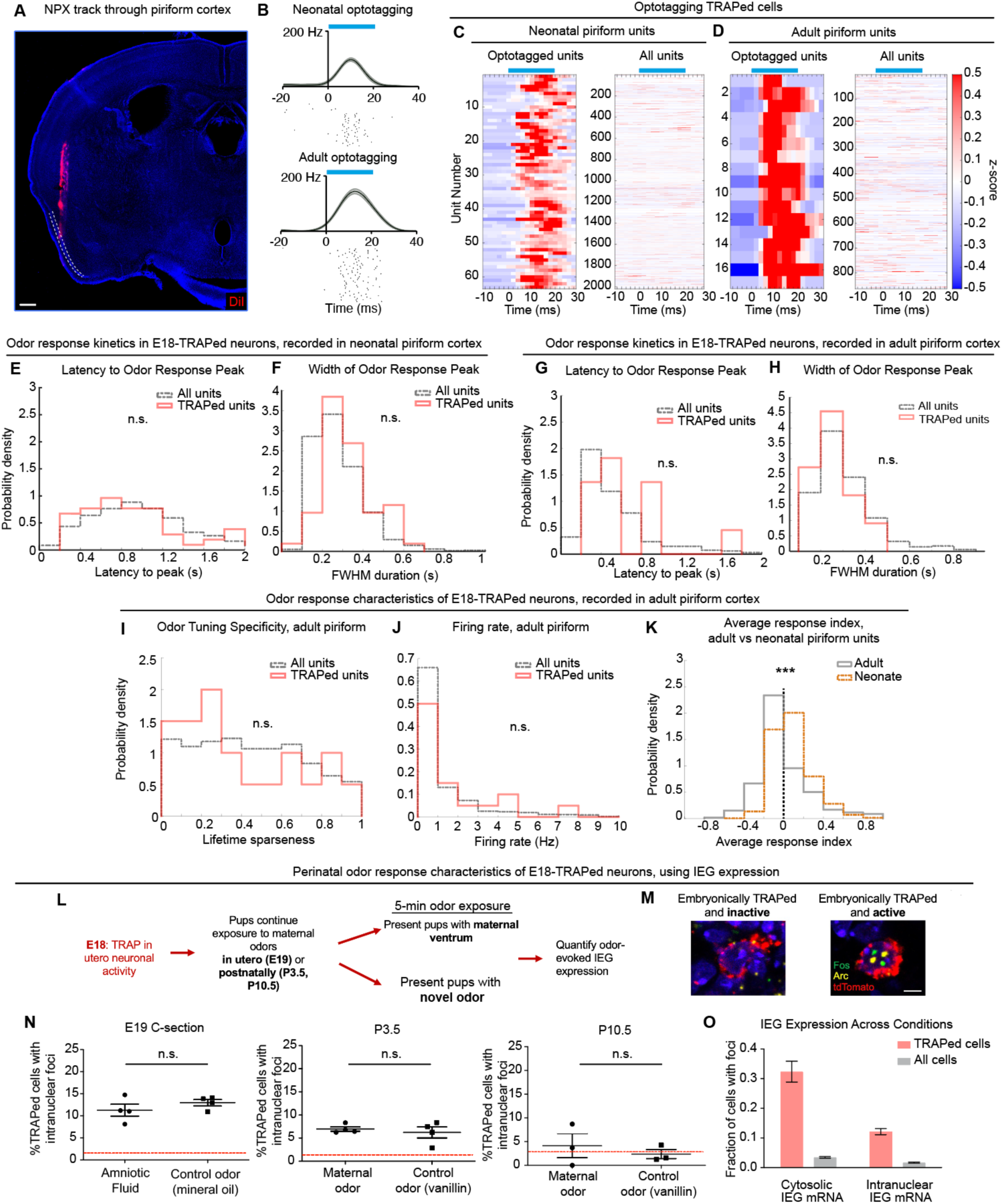
Optotagging E18-TRAPed units, further odor response characteristics of E18-TRAPed neurons in neonatal and adult piriform cortex *in vivo*, and perinatal odor response of E18-TRAPed neurons using IEG expression, related to Figure 4. (A) Example histological section showing DiI-stained Neuropixels track entering piriform cortex (dotted lines outline layer 2 of piriform cortex), demonstrating that Neuropixels spans all piriform cortical layers. Scale bar, 400 μm. (B) PSTHs and spike trains of light-responsive units in neonatal (upper) and adult (lower) piriform of E18-TRAPed mice. Blue bar indicates light presentation. (C, D) Summary of optotagged TRAPed cell z-scored during optotagging compared to all units, recorded in neonatal piriform (C) and adult piriform (D) corresponding to data in Figure 4. (E, F) Latency to peak activity during odor presentation (E) and width of peak activity (F; calculated as full-width half-max of peak) in TRAPed vs all units recorded in neonatal piriform. Latency and width of odor response peaks are only shown for individual cell–odor pairs that have significant responses (response index > 0.4). (G, H) Latency to peak activity during odor presentation (E) and width of peak activity (F; calculated as full-width half-max of peak) in TRAPed vs all units recorded in adult piriform. Latency and width of odor response peaks are only shown for individual cell–odor pairs that have significant responses (response index > 0.4). (I, J) Lifetime sparseness (I; See Methods) and baseline firing rate (J) of E18-TRAPed vs all neurons recorded in adult piriform cortex. Latency and width of odor response peaks are only shown for individual cell-odor pairs that have significant responses (response index > 0.4). n_TRAP_ = 17 units, n_Total_ = 864 units, recorded across 4 adult mice; n.s., not significant, t-test with bootstrapping. (K) Average response indices of all piriform units in neonate vs adult (n_Neo_ = 2020, n_Adult_ = 864; ***p<0.001, t-test with bootstrapping). (L) Experimental design for reactivation of E18-TRAPed neurons following brief exposure to a maternal or novel odor. IEG, immediate early gene. (M) Example RNAscope images of E18-TRAPed neurons that are inactive (left) or active and containing intranuclear IEG transcripts (right). Scale bar, 3 μm. (N) Summary of activation of TRAPed neuron following exposure to maternal or novel odor at E19, P3.5 or P10.5. Each dot represents percentage of TRAPed neurons that contained intranuclear IEG transcripts. n = 50 cells per animal, n = 4 animals per condition per age; short lines represent 25-75^th^ percentile, long line represents mean; red dotted lines correspond to percentage of all cells in imaged regions of piriform cortex that contained intranuclear IEG transcripts. (O) Quantification of IEG transcript expression in TRAPed vs all cells across all ages and odor stimulation conditions. Error bars, SEM.

**Figure S5.**
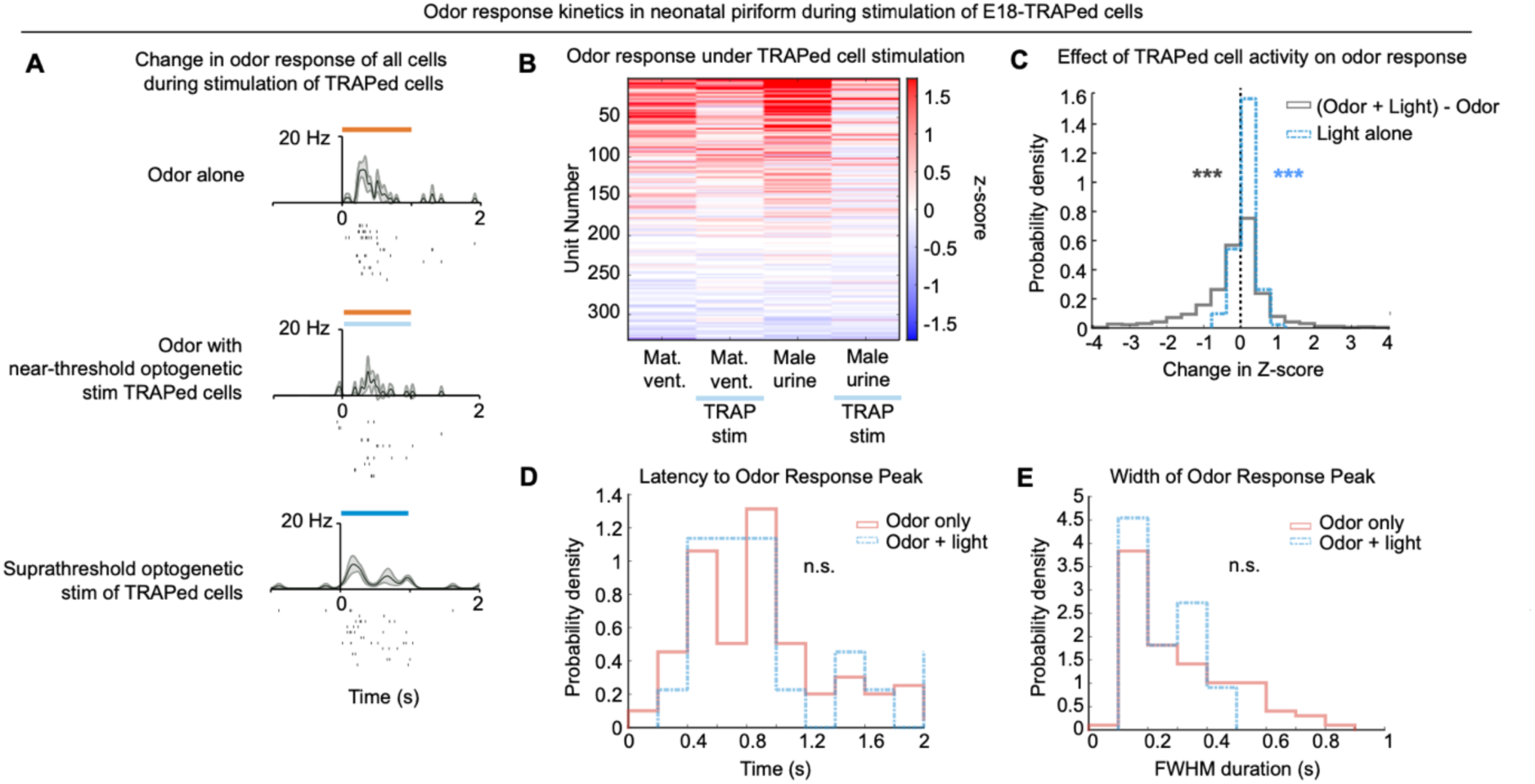
Effect of near-threshold optogenetic stimulation of E18-TRAPed cells on odor response in neonatal piriform cortex. (A) Example PSTHs of nonTRAPed unit in neonatal piriform in response to odor presentation (upper), odor presentation with near-threshold stimulation (see **Methods**) of E18-TRAPed cells (middle), or suprathreshold stimulation of E18-TRAPed cells (lower). Orange bar indicates odor presentation, blue bar indicates light presentation. (B) Odor response z-score of neonatal piriform units to two tested odorants with and without near-threshold optogenetic stimulation of TRAPed cells during odor presentation. Cells are sorted by average z-score in response to all four odor presentations (n = 332 units; blue bar indicates light presentation during odor; Mat. Vent.: maternal ventrum). (C) Distributions of z-score changes of neonatal piriform units in response to near-threshold optogenetic stimulation of TRAPed cells. Gray trace shows the difference in z-score (calculated as response during odor presentation compared to baseline activity) between odor presentation with TRAPed cell stimulation and odor presentation alone. Positive values along the x-axis suggests an increase in odor response when TRAPed cells are stimulated during odor presentation. The majority of the gray distribution lies to the left of 0 (mean = –0.273, ***p<0.001 in gray text, one-sample t-test), suggesting a decrease in odor response when TRAPed cells are stimulated during odor presentation. Blue trace shows the z-score changes with near-threshold optogenetic stimulation of TRAPed cells in the absence of odor presentation. The blue distribution lies slightly to the right of 0 (mean = 0.067, ***p<0.001 in blue text, one-sample t-test), suggesting that optogenetic stimulation TRAPed cells at baseline leads to a mild increase in activity of the recorded population (n = 332 units). (D, E) Latency (D) and width (E) of odor responses of neonatal piriform units with (blue trace) and without (orange trace) at-threshold optogenetic stimulation of E18-TRAPed cells during odor presentation. Latency and width of odor response peaks are only shown for individual cell– odor pairs that have significant responses (response index > 0.4) (n.s., not significant; t-test with bootstrapping).

**Figure S6.**
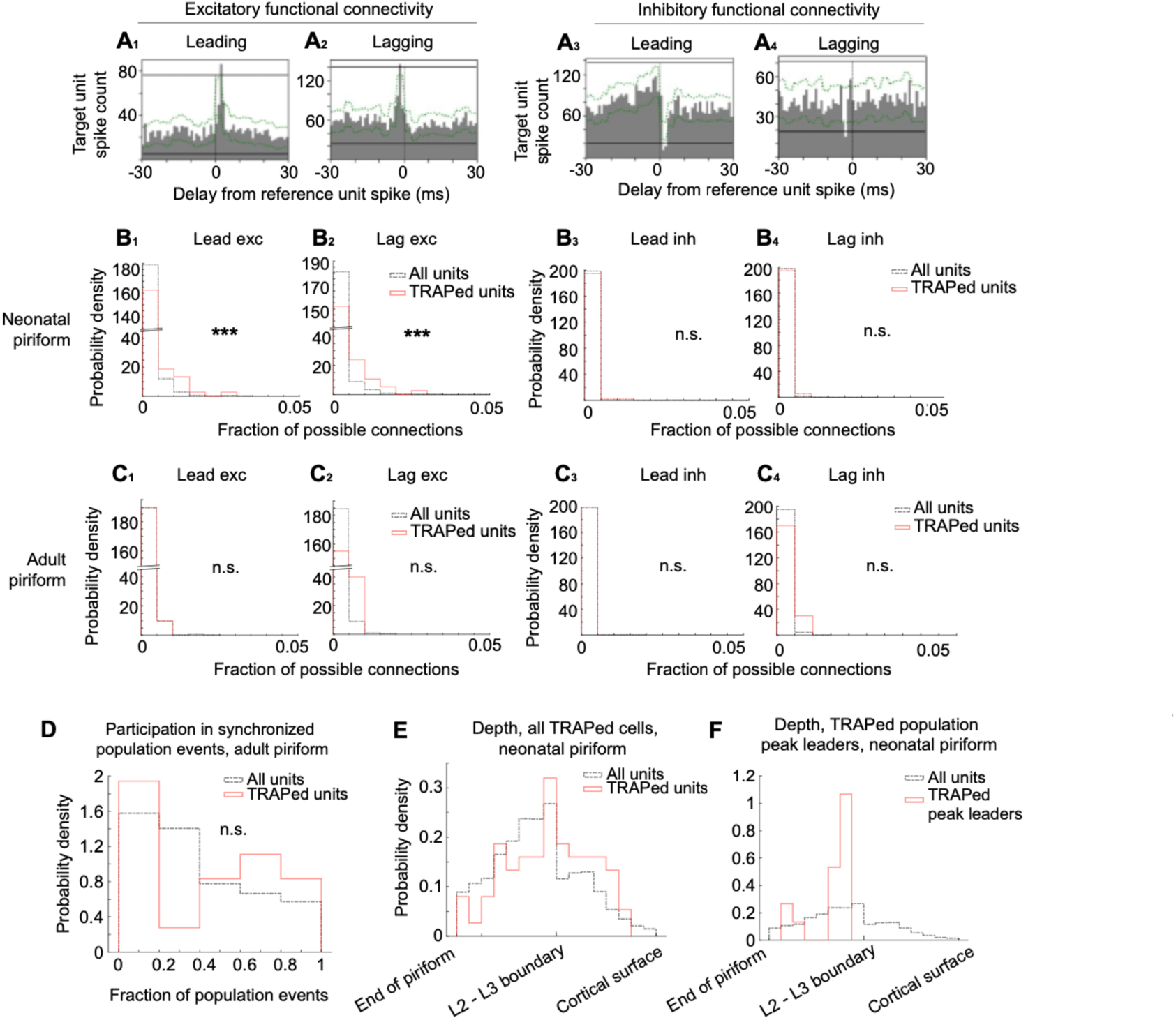
Functional connectivity and anatomical characteristics of E18-TRAPed piriform neurons *in vivo*, and additional data on E18-TRAPed cell activity in adult piriform cortex *in vivo*, related to Figure 5. (A) Example cross-correlograms of pair of units with excitatory leading (A_1_) or lagging (A_2_) or inhibitory leading (A_3_), or lagging (A_4_) functional connectivity. Green dotted traces represent significance envelope calculated via jitter method^79^. Solid horizontal black lines within the plot indicate the upper and lower boundaries of significance. (B) Distribution of frequency of each subtype of functional connectivity in E18-TRAPed vs all units in neonatal piriform cortex. **p<0.005; n.s., not significant; t-test with bootstrapping. (C) Distribution of frequency of each subtype of functional connectivity in E18-TRAPed vs all units in adult piriform cortex. n.s., not significant; t-test with bootstrapping. (D) Fraction of synchronized population events in which E18-TRAPed vs all units participated, recorded in adult mice. (E, F) Distribution of depth indices of TRAPed (E) or TRAPed population peak leaders (F) vs all units in neonatal piriform. n_TRAP,neonate_ = 26, n_Total,neonate_ = 564, n_TRAP,adult_ = 17, n_Total,adult_ = 864 across 5 neonatal mice and 4 adult mice. x-axes are from dorsal to ventral. Note that TRAPed peak leader units are more dorsal (deeper) than L2-L3 boundary and are thus located in L3.

**Figure S7.**
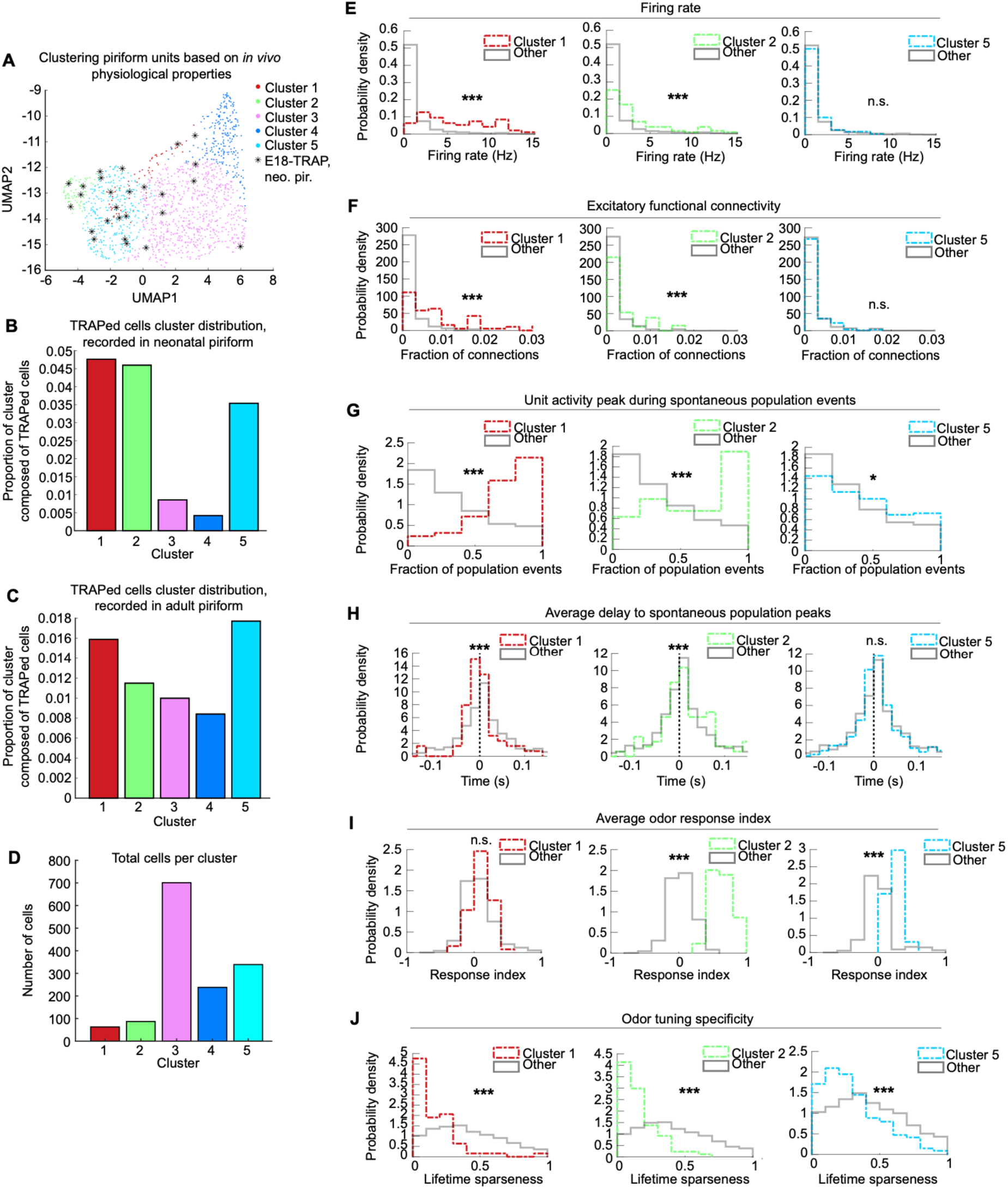
TRAPed cell distribution across piriform unit types clustered using *in vivo* anatomical and physiological features, related to Figure 5. (A) UMAP representation of units grouped by k-means clustering of *in vivo* anatomical and physiological properties (see methods for list of features). n = 1428 total units across 9 mice in which all features were recorded, n_TRAP_ = 26 units; neo. pir., neonatal piriform. (B, C) Proportion of each cluster that is composed of E18-TRAPed cells recorded in neonatal piriform cortex (B; shown as asterisks in A) or adult piriform cortex (C; not highlighted in A). (D) Total number of units per cluster from (A). (E–J) Distribution of firing rate (E), excitatory functional connectivity (F), unit activity peak co-occurring with spontaneous population activity peak (G), and average delay to spontaneous population peak (H), average odor index (I), and lifetime sparseness (J), for the three clusters in which E18-TRAPed cells had the most representation. Distributions for clusters of interest at shown in colored traces, compared to units in all other clusters in grey traces. *p < 0.00033, ***p< 0.0000033, n.s. not significant; bootstrapped t-test with Bonferroni multiple-comparisons correction.

**Figure S8.**
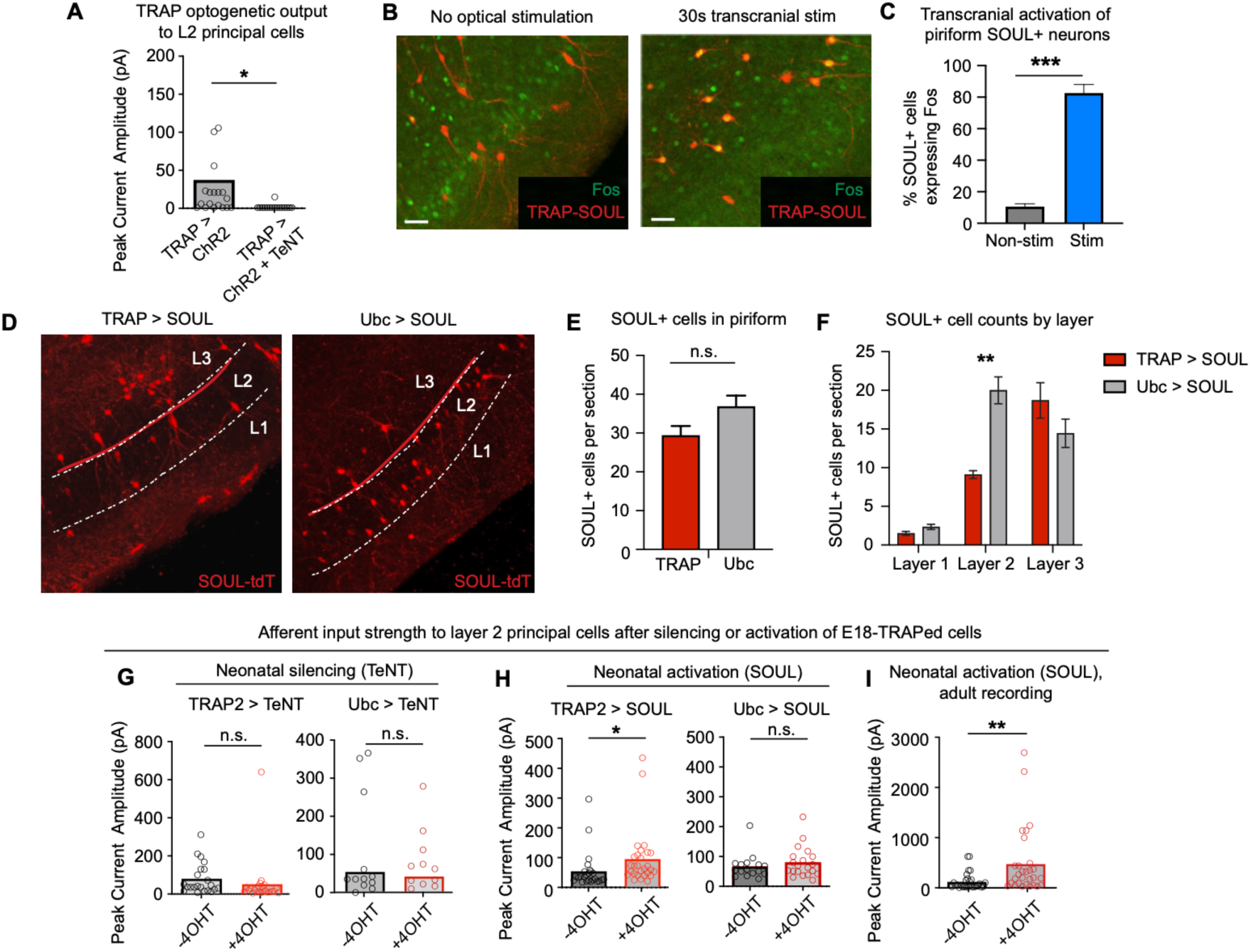
Further characterization of activity manipulation experiments, related to **Figure 8**. (A) Quantification of optogenetic input from TRAPed cells to nonTRAPed layer 2 principal neurons in animals with TRAPed inputs intact (TRAP > ChR2) and TRAPed outputs blocked (TRAP > ChR2 + TeNT). n_ChR2_ = 20 cells, n_ChR2+TeNT_ = 16 cells; *p < 0.05, unpaired t-test (B, C) Example (B) and quantification (C) of Fos expression in SOUL+ cells evoked by brief transcranial optical stimulation in a neonatal *TRAP2;SOUL* animal (n = 3 animals per condition, ***p<0.0005, unpaired t-test). Error bars, SEM. Scale bar, 40 μm. (D–F) Example (D) and quantification of SOUL-expressing cells in *TRAP2;LSL-SOUL* (TRAP > SOUL) vs *Ubc-CreER^T2^;LSL-SOUL* (Ubc > SOUL), by total count in piriform (E) and distribution by layer (F). n = 3 per condition, **p<0.005, n.s., not significant; two-way ANOVA. Scale bar, 50 µm. *LSL, loxP-stop-loxP*. (G) Plot of peak current amplitude of afferent inputs in *TRAP2;LSL-TeNT* (TRAP > TeNT; *LSL*, *loxP-stop-loxP*) or *Ubc-CreER^T2^;LSL-TeNT* (Ubc > TeNT) animals compared to fostered littermates that did not receive 4OHT. n_-4OHT,TRAP_ = 22 cells across 5 mice, n_+4OHT,TRAP_ = 23 cells across 6 mice, n_-4OHT,Ubc_ = 11 cells across 3 mice, n_+4OHT,Ubc_ = 12 cells across 3 mice; *p<0.05, n.s. not significant; unpaired t-test. (H) Plot of peak current amplitude of afferent inputs in stimulated *TRAP2;SOUL* (TRAP > SOUL) or *Ubc-CreER^T2^;SOUL* (Ubc > SOUL) animals compared to fostered littermates that did not receive 4OHT. n_-4OHT,TRAP_ = 25 cells across 5 mice, n_+4OHT,TRAP_ = 28 cells across 3 mice, n_-4OHT,Ubc_ = 13 cells across 2 mice, n_+4OHT,Ubc_ = 17 cells across 3 mice; *p<0.05, n.s. not significant; unpaired t-test. (I) Plot of peak current amplitude of afferent inputs in adult *TRAP2;SOUL* (TRAP > SOUL) mice animals compared to fostered littermates that did not receive 4OHT following neonatal activation of E18-TRAPed cells. n_-4OHT,TRAP_ = 23 cells across 4 mice, n_+4OHT,TRAP_ = 24 cells across 4 mice; *p<0.05, **p<0.005, n.s. not significant; unpaired t-test.

